# Beyond spike-timing-dependent plasticity: a computational study of plasticity gradients across basal dendrites

**DOI:** 10.1101/063719

**Authors:** Jacopo Bono, Claudia Clopath

**Author notes:** Correspondence and requests for materials should be addressed to C.C.

## Abstract

Synaptic plasticity is thought to be the principal mechanism underlying learning in the brain. Models of plastic networks typically combine point neurons with spike-timing-dependent plasticity (STDP) as the learning rule. However, a point neuron does not capture the complexity of dendrites, which allow non-linear local processing of the synaptic inputs. Furthermore, experimental evidence suggests that STDP is not the only learning rule available to neurons. Implementing biophysically realistic neuron models, we studied how dendrites allow for multiple synaptic plasticity mechanisms to coexist in a single cell. In these models, we compared the conditions for STDP and for the synaptic strengthening by local dendritic spikes. We further explored how the connectivity between two cells is affected by these plasticity rules and the synaptic distributions. Finally, we show how memory retention in associative learning can be prolonged in networks of neurons with dendrites.

## INTRODUCTION

Our brain continuously processes and stores novel information, providing us with a remarkable flexibility to adapt and learn in a continuously changing environment. Understanding which mechanisms underlie this plasticity is an important step towards understanding how sensory experience is processed in cortical areas. A seminal idea, now widely known as Hebb’s postulate, suggested that synapses are the neurological substrates for learning [Hebb, 1949, Berlucchi and Buchtel, 2009]. In short, Hebb proposed that a neuron A persistently taking part in firing a neuron B, leads to increased synaptic efficacy from neuron A to B. Experiments later showed that high-frequency stimulation indeed evoked long-lasting increased efficacy, termed long-term potentiation (LTP), at hippocampal synapses [Bliss and Lømo, 1973]. Long-term depression (LTD), predicted as a mechanism to balance LTP [Stent, 1973, Bienenstock et al., 1982], was later shown to be evoked by low-frequency stimulation [Dudek and Bear, 1992].

A more complex picture of synaptic plasticity emerged, uncovering how the precise timing between the presynaptic and postsynaptic activity influences plasticity. When the presynaptic neuron fires just before the postsynaptic neuron, the synapse is potentiated. Instead, reversing the firing order leads to synaptic depression [Gerstner et al., 1996, Markram, 1997, Bi and Poo, 1998]. This form of plasticity, where the precise timing of spikes determines the subsequent synaptic changes, is named spike-timing-dependent plasticity (STDP) and is currently a widely used learning rule in computational studies [Caporale and Dan, 2008, Morrison et al., 2008, Feldman, 2012].

Furthermore, the synaptic changes were shown to be additionally dependent on the rate of presynaptic and postsynaptic activity [Markram, 1997, Sjöström et al., 2001], postsynaptic depolarisation [Artola et al., 1990, Ngezahayo et al., 2000, Harvey and Svoboda, 2007], multiple pre-or postsynaptic spikes [Froemke and Dan, 2002, Wang et al., 2005, Nevian and Sakmann, 2006] and dendritic location of the synapse [Froemke et al., 2005, Sjöström and Häusser, 2006, Letzkus et al., 2006, Froemke et al., 2010]. These results have led to refined models for synaptic plasticity, among which are mechanistic models based on the calcium hypothesis [Shouval et al., 2002, Graupner and Brunel, 2012], triplet models [Pfister and Gerstner, 2006] and a voltage-dependent phenomenological model [Clopath et al., 2010] which we use in this study.

The STDP mechanism relies on the generation of action potentials and their subsequent backpropagation into the dendrites. These two requirements limit the capacity of STDP in the following cases. Firstly, canonical pair-based STDP cannot account for activity-dependent learning with weak inputs, which are not powerful enough to evoke action potentials. Secondly, the attenuation or failure of the back-propagating action potential (bAP) provides a problem for STDP in the dendritic regions far from the soma [Froemke et al., 2005, Sjöström and Häusser, 2006, Letzkus et al., 2006]. Furthermore, an increasing number of experimental studies have revealed plasticity mechanisms which do not rely on postsynaptic action potential generation, but instead on local postsynaptic dendritic spikes [Golding et al., 2002, Kampa et al., 2006, Gordon et al., 2006, Gambino et al., 2014, Brandalise and Gerber, 2014, Kim et al., 2015, Cichon and Gan, 2015] or subthreshold events for dendritic spikes [Weber et al., 2016, Sandler et al., 2016]. The power of the STDP mechanism has since been highly debated [Lisman and Spruston, 2005, Hardie and Spruston, 2009, Lisman and Spruston, 2010, Schulz, 2010, Shouval et al., 2010, Frégnac et al., 2010, Buchanan and Mellor, 2010].

Taken together, these considerations indicate that pair-based STDP forms only part of the plasticity-machinery a neuron offers. Since the learning rules shape the connections in our brain, which ultimately affect the processing of information, it is crucial to understand which forms of plasticity are most prominent in neurons. In this article, we investigated how dendrites allow for multiple plasticity rules in a single cell and attempt to compare their relative importance. In biophysical models of a cortical layer 5 and layer 2/3 pyramidal neuron, we implemented the voltage-based STDP rule (vSTDP, [Clopath et al., 2010]) which reproduces the various plasticity mechanisms described above [Artola et al., 1990, Markram, 1997, Ngezahayo et al., 2000, Sjöström et al., 2001, Froemke and Dan, 2002, Golding et al., 2002, Wang et al., 2005, Froemke et al., 2005, Sjöström and Häusser, 2006, Nevian and Sakmann, 2006, Letzkus et al., 2006, Harvey and Svoboda, 2007, Froemke et al., 2010, Gambino et al., 2014], and we focussed on the basal dendritic tree. Our simulations reproduce experimental evidence of a gradient of plasticity along these dendrites, with STDP dominating proximally and a dendritic-spike-dependent LTP (dLTP) distally. We then investigated how synaptic distribution affects the connectivity between two neurons and how NMDA spikes can determine the sign of plasticity at other synapses. Finally, we explored the influence of dLTP in the context of associative learning in single cells and networks of simplified neurons, in particular, we found that dLTP can prolong memory retention.

## RESULTS

We modelled a cortical layer 5 and layer 2/3 pyramidal neuron. The reconstructed morphologies of the neurons were taken from [Acker and Antic, 2009] and [Branco et al., 2010] respectively. In the main text we will only discuss results of the layer 5 model (Figure 1a), however we refer to the Supplementary Figures S1 and S2 for similar results using the layer 2/3 neuron. In our morphologically reconstructed neurons, we added Na^+^, K^+^ and Ca^2+^ conductances based on the fit to experimental data performed in [Acker and Antic, 2009] (see methods). All simulations of the biophysical neurons were performed using Python with the Brian2 simulator [Goodman, 2008]. The synapses have both AMPA and NMDA components as in [Branco et al., 2010]. Our model was able to reproduce the observed integration gradient along thin basal dendrites as reported in Branco and Häusser both experimentally and in a computational model [Branco and Häusser, 2011] (Figure 1a,b). This gradient was shown to arise from a difference in input impedance along the thin dendrites, resulting in greater depolarisation by single excitatory postsynaptic potentials (EPSPs) at distal locations on the dendrite compared to proximal locations. In turn, the higher depolarisation activates the supra-linear NMDA current. In our layer 5 model, 16 out of 17 terminal compartments evoked NMDA spikes while reproducing the stimulation protocol in [Branco and Häusser, 2011] (Figure 1a,b), while in the layer 2/3 model 10 out of 15 terminal compartments evoked NMDA spikes (Supplementary Figure S1a,b). In our simulations, when choosing distal compartments we used only those that evoked NMDA spikes, unless otherwise mentioned. To avoid further ambiguities in light of current literature, we point out to the reader that with *distal* we will denote the regions furthest from the soma on *basal* dendrites, as opposed to the apical tuft dendrites, and with *proximal* we mean the regions closer to the soma on *basal* dendrites, as opposed to the whole basal dendritic tree.

**Figure 1.**
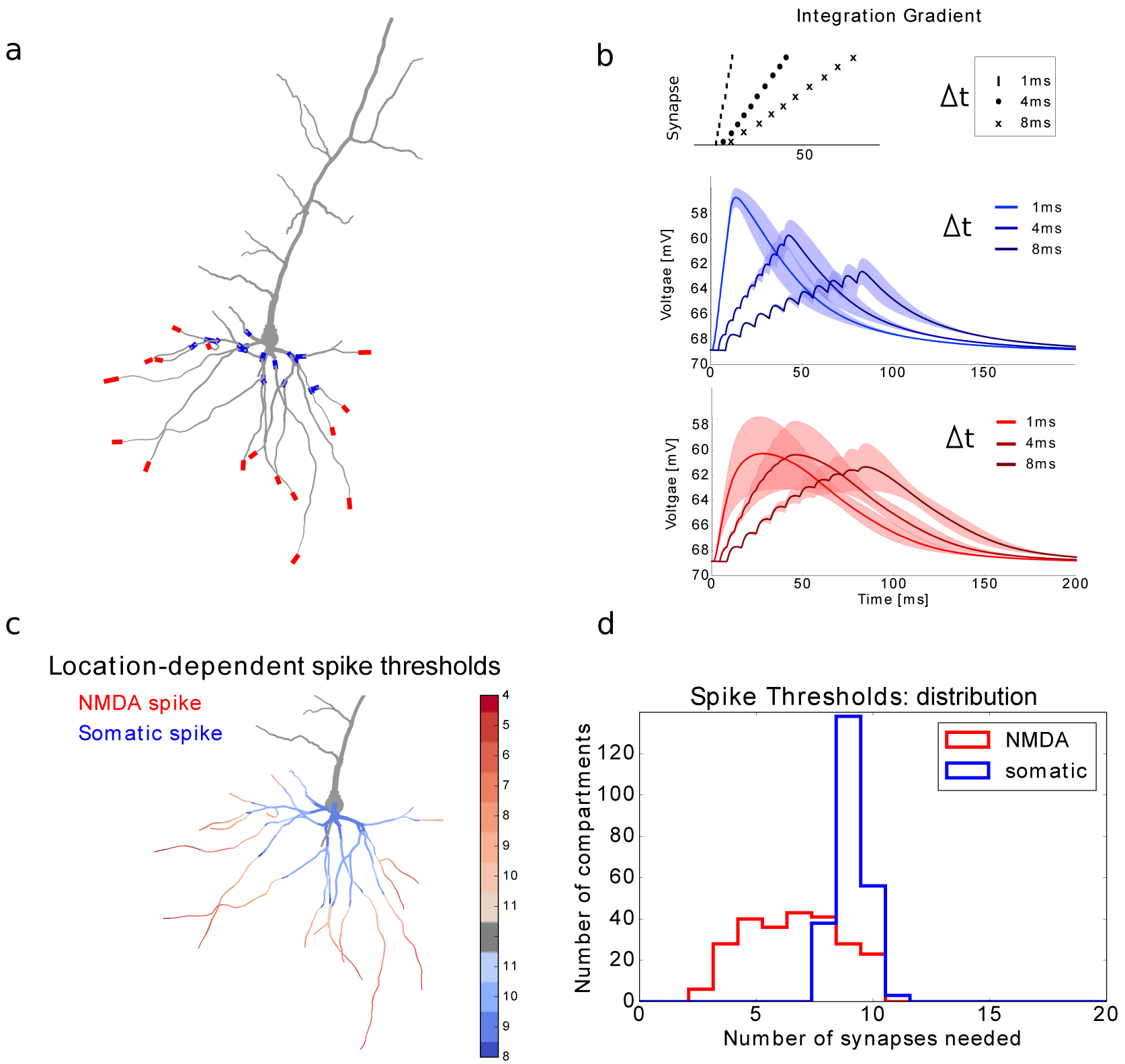
Location dependence of local and global regenerative events. (a) Proximal (blue) and distal (red) location on a thin basal branch of the detailed neuron model. (b) Temporal integration gradient along a basal dendrite (as observed in [Branco and Häusser, 2011]). Top panel: activation sequences with different interspike intervals. Middle panel: proximal stimulation requires high temporal coincidence, as the somatic depolarisation drops substantially when inputs are not active synchronously. Bottom panel: distal stimulation results in similar somatic depolarisation for a bigger range of interspike intervals. (c) For each dendritic compartment in the neuron model, synapses are activated until either an NMDA spike or an action potential is generated. Stimulating proximal compartments results in an action potential before reaching the threshold for NMDA spikes (blue). Stimulating distal regions results in NMDA spike generation while staying subthreshold for somatic spikes (red). Intenser colors denote that less synapses are needed to evoke a spike. The numbers next to the colorbar represent the amount of synapses needed for a spike. (d) For each compartment in (c), the number of synapses needed to evoke the spikes was stored and represented as a histogram. NMDA spikes (red) generally require around 5 activated synapses in our model, while somatic spikes (blue) require around 10 synapses.

For the network simulations, we developed a reduced neuron model where each dendrite consists only of two compartments (see methods) and fitted it to the biophysical model (Supplementary Figure S4). The reduced model was developed in Python, the code for both models and all simulations is available on ModelDB (https://senselab.med.yale.edu/modeldb/).

### 1/ Location dependence of local and global regenerative events

Dendrites allow for various local regenerative events, named dendritic spikes [Major et al., 2013]. Dendritic spikes in basal dendrites of cortical pyramidal neurons are usually evoked by the activation of NMDA receptor mediated channels, and called NMDA spikes [Gordon et al., 2006, Major et al., 2008, Antic et al., 2010]. We firstly quantify the NMDA-dependent integration gradient in our biophysical model in more detail: for each of the dendritic compartments, we recorded which spike (somatic or NMDA spike) requires the least synaptic input and we also stored the amount of synapses needed. In our model, synapses clustered on the proximal dendritic compartments will result in the firing of a somatic spike before generating an NMDA spike, while synapses clustered on the distal dendritic compartments evoke an NMDA spike without generating a somatic spike (Figure 1c). Moreover, the number of synapses needed to generate an NMDA spike is generally much lower than that for a somatic spike (Figure 1d). Our model also confirms that increasing the synaptic input on a distal compartment usually will not result in a combination of NMDA spike and somatic spike: increased input on such compartment will increase the duration of the plateau, but not the amplitude, resulting in a longer (but still subthreshold) signal at the soma [Major et al., 2008]. A combination of the NMDA spike and other inputs arriving at different compartments, however, may be especially useful for evoking somatic spikes, as will be discussed in section 4.

### 2/ Location dependence of STDP and dLTP

We wondered how the integration gradient described in the previous section affects synaptic plasticity. For this purpose, a local voltage-dependent STDP rule (vSTDP) was implemented in our model, based on [Clopath et al., 2010]. The parameters were fitted as to reproduce the depolarisation- and rate-dependence of plasticity [Artola et al., 1990, Ngezahayo et al., 2000, Markram, 1997, Sjöström et al., 2001, Harvey and Svoboda, 2007] (Figure 2b). Our model is consistent with experimental results reporting a reduced potentiation window for increasing distances from the soma, due to the attenuation of the back-propagating action potential (bAP) [Froemke et al., 2005, Sjöström and Häusser, 2006, Letzkus et al., 2006] (Figure 2c). We also found that the same plasticity rule was able to account for local, dendritic-spike-dependent LTP, observed in vitro [Golding et al., 2002, Gordon et al., 2006, Kampa et al., 2006, Brandalise and Gerber, 2014, Weber et al., 2016] and in vivo [Gambino et al., 2014, Cichon and Gan, 2015]. Since vSTDP depends on voltage rather than postsynaptic spiking, a large postsynaptic depolarisation caused by a dendritic spike is sufficient for LTP and no back-propagating action potential (bAP) is required (Figure 2e).

**Figure 2.**
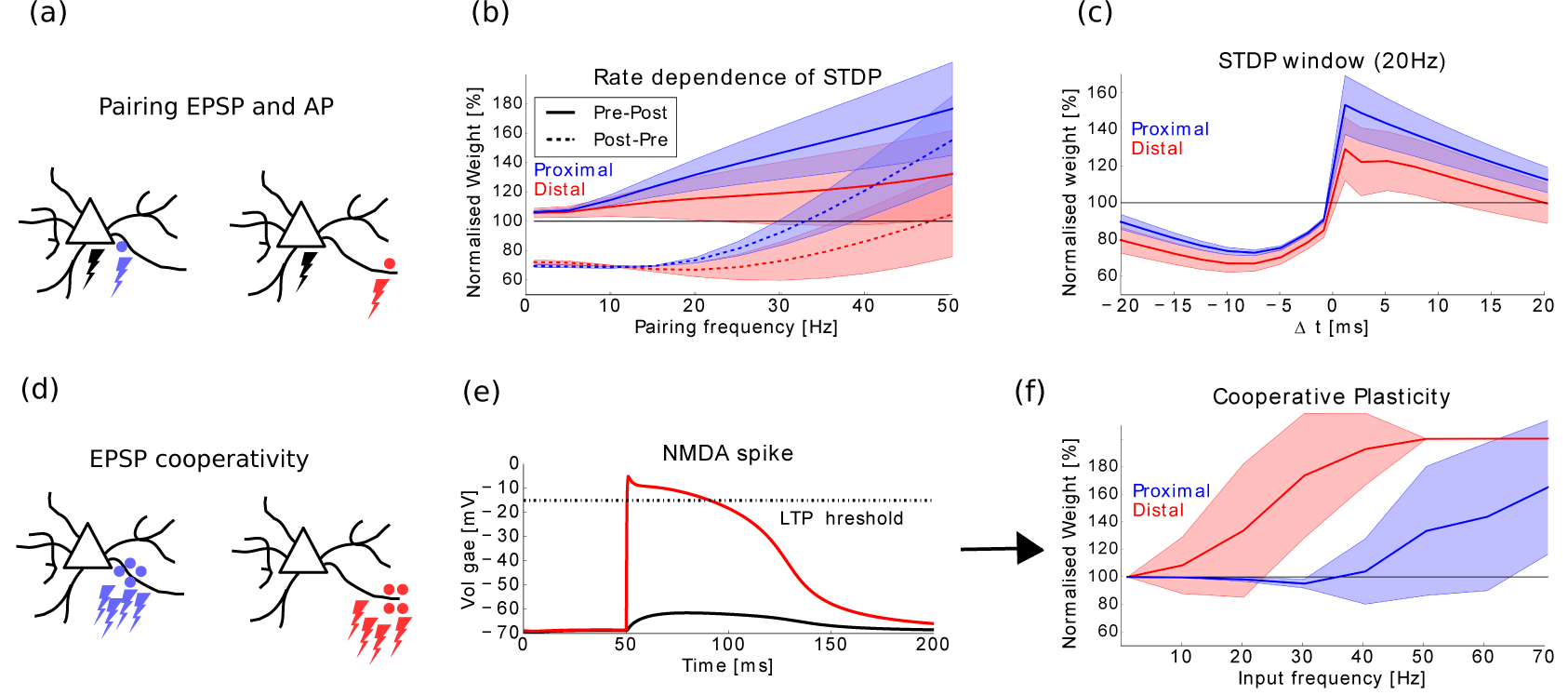
Plasticity gradient along basal dendrites. (a) Action potentials (black lightning) is paired with either proximal (blue) or distal (red) synaptic activations. (b) The pairing protocol as in [Sjöström et al., 2001] is simulated. Action potentials and EPSPs are paired with a fixed interval of 10ms, either pre-post (full lines) or post-pre (dashed lines). The pairings are repeated 15 · 5 times at a fixed frequency (see methods). The protocol is repeated for several pairing frequencies (horizontal axis) and the normalised weight change is displayed (vertical axis). The parameters of the plasticity model were chosen to qualitatively reproduce the experimentally observed rate-dependence, where low rates lead to depression only, intermediate rates allow for both potentiation and depression depending on the spike timings, and high rates lead to potentiation only. Distal synapses show less potentiation due to the attenuation of the bAP. (c) The same protocol as in (b) is repeated at a pairing frequency of 20 Hz, but the interspike interval is now varied (horizontal axis). The normalised weight change (vertical axis) shows the standard STDP window, with a reduced window for potentiation at distal synapses due to the attenuation of the bAP. (d) A group of synapses on a proximal compartment (blue) or distal compartment (red) are activated by a Poisson process. (e) Distal groups of synapses can elicit NMDA spikes (red trace), which cross the LTP threshold of the plasticity model and can therefore result in potentiation while remaining subthreshold in the soma (black trace). (f) When activating the synaptic groups as in (d), LTP is observed at much lower average activation rates for the distal group (red trace) compared to the proximal group (blue trace). This local, dLTP by NMDA spikes compensates for the lack of potentiation by STDP (b,c).

Since NMDA spikes are more easily evoked at the distal regions of a dendrite (Figure 1c,d), we investigated whether dLTP, without the need for action potentials, is more easily evoked at distal locations. For this purpose, we stimulated 10 synapses either placed distally or proximally on the same dendrite. The synapses were all activated using a Poisson process with the same average rate. The synaptic group at the distal part of the dendrite led to potentiation for substantially lower activation rates as opposed to the proximal group (Figure 2f).

Our simulations are consistent with the experimental observation of a plasticity gradient along basal dendrites: STDP is more effective in proximal regions, while dLTP is more easily evoked at distal regions. The latter mechanism could rescue the distally reduced potentiation caused by an attenuated bAP. Note that so far we have only discussed an additional mechanism for LTP at distal locations, evoked by NMDA spikes. However, the depression at distal locations can occur as well simply by inputs that remain sub-threshold for NMDA spikes. This can be achieved by asynchronous or low-rate activations of distal synapses (see Figure 2f), or by providing simultaneous inhibitory input.

### 3/ Synapse distribution influences connectivity

We wondered how the plasticity mechanisms described in the previous paragraph influence the connectivity between two neurons. For this purpose, we simulated two biophysical neurons that are mutually connected. We varied the distribution of synapses in the following ways: they are either randomly chosen across the basal tree (Figure 3a-d), randomly distributed across only proximal compartments or only distal compartments (Figure 3e-h), or grouped onto a single proximal or a single distal compartment (Figure 3i-l and o-r). Furthermore, we activated the neurons by either generating Poisson spike trains over a range of rates (Figure 3a-n), or by activating one neuron always before the second (Figure 3o-r).

**Figure 3.**
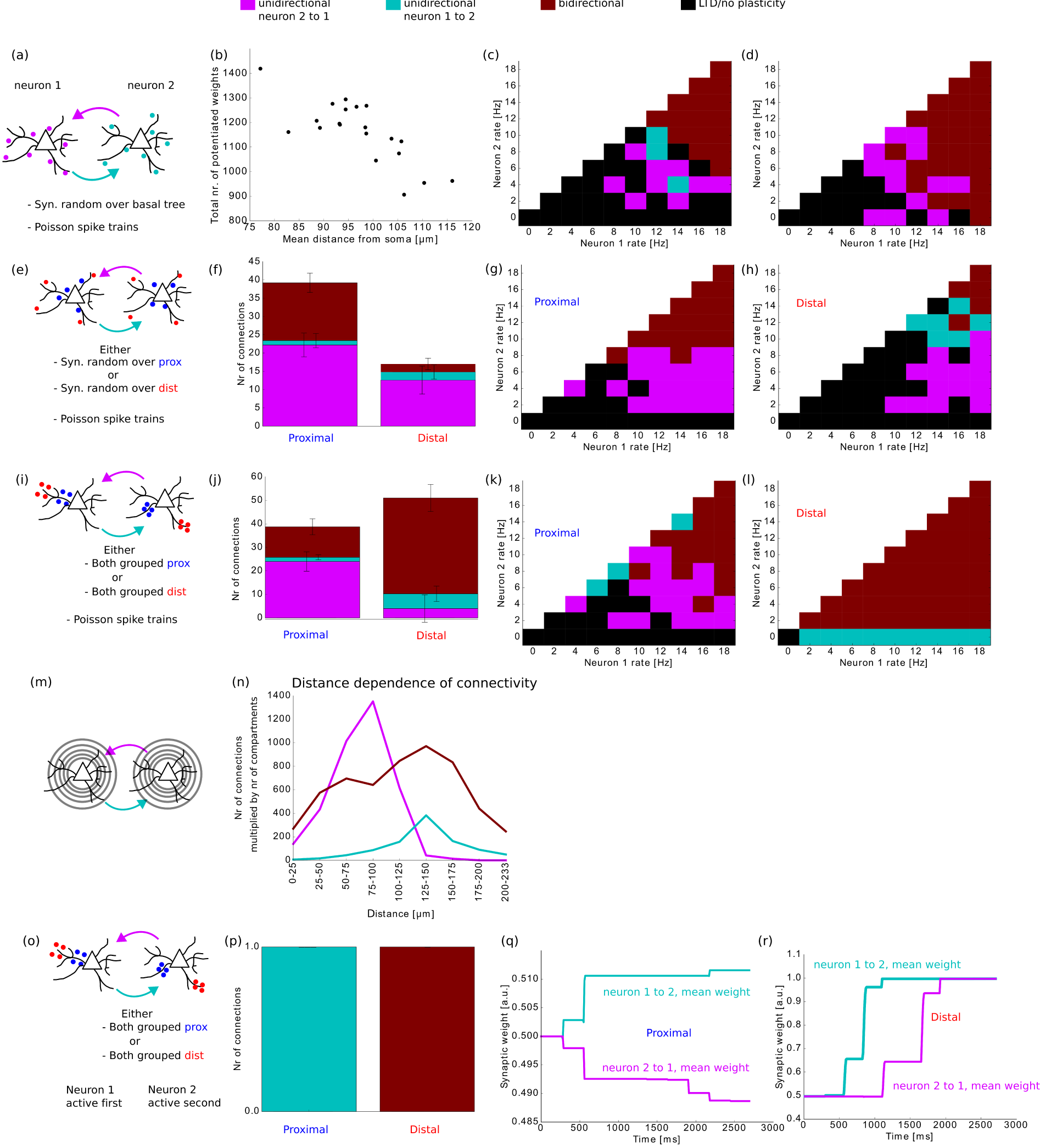
Predicted connectivity along dendrites. (a) Two neurons are mutually connected: the synapses are randomly distributed across the basal dendrites and both neurons are activated generating Poisson spike trains. (b) The mean distance of all synapses to the soma is inversely correlated to the number of strengthened synapses (Pearson r=−0.785, p=4e-5). (c,d) Two examples of final connectivity. Color codes can be found at the legend on top of the figure. (e) Same as (a), but the synapses are now constraint to either proximal or distal compartments. (f) More synapses are strengthened if targeting proximal regions when compared to distal synapses. (g,h) Examples of final connectivity for the proximally distributed case (g) and the distal distributed case (h). (i) Analogous as (e), but synapses are now clustered onto one proximal or one distal compartment. (j) By comparing with (f), proximal synapses are less influenced by the clustering of synapses. Clustering of distal synapses leads more easily to LTP and therefore bidirectional connections are dominant. (k,l) Examples of final connectivity for the proximally clustered case (k) and the distally clustered case (l). (m) We divide our basal dendritic tree in regions depending on the distance from the soma. For each distance-range, synapses are either randomly distributed among the relevant compartments, or clustered on one of the compartments. (n) Mean results of connectivities for the distributed and the clustered simulations in each distance-range are summed. The values are multiplied by the number of compartments in the respective distance-range. (o) A temporal order is imposed: neuron 1 is always activated before neuron 2. The synapses are clustered either proximally or distally. (p) In the proximal case, all simulations result in unidirectional connections from neuron 1 to neuron 2. In the distal case, only bidirectional connections are observed. (q,r) Example of the evolution of the mean synaptic weights during the simulation for the proximal case (q) or the distal case (r).

We then investigated the synaptic changes caused by these different synaptic distributions and stimulation protocols. We first used the completely random distribution of 20 synapses (Figure 3a) while evoking Poisson spike trains in both neurons. For firing rates lower than 12Hz, we mainly observed synaptic depression or unidirectional connections from the neuron with lowest firing rate towards the neuron with higher firing rate. For higher rates, bidirectional connections were formed (Figure 3b,c). Since we were interested to see whether the distance from the soma affected the development of strong connections, we calculated the average distance of synapses from the soma in each simulation and plotted this as a function of the total number of potentiated synapses (Figure 3d). The data has a strong negative correlation coefficient (−0.785) suggesting that proximity to the soma favours LTP. To further investigate this, we now repeated the simulation but while distributing 10 synapses either only over the proximal or only the distal compartments (Figure 3e, the compartments used in these simulations are the same as shown in Figure 1a). The proximally connected neurons indeed clearly favour potentiation compared to the distally connected neurons, resulting in unidirectional or bidirectional connectivity depending on the firing rates (Figure 3f,g,h). We can understand the results as follows: since the synapses in the previous simulations where randomly distributed, the probability to find groups of synapses connected to the same compartment is negligible. In this case, NMDA spikes will not be evoked and STDP will dominate. The height of the bAP is therefore crucial and favours proximal synapses.

We then changed the distribution of synaptic connections again and clustered all synapses together onto one compartment, either proximally or distally (Figure 3i). When the neurons are proximally connected in this way, the connectivity is not substantially different from the randomly distributed synaptic configurations (compare Figures 3j,k with Figures 3f,g). However, for distal locations an NMDA spike can easily be evoked and in turn this changes the plasticity substantially (compare Figures 3j,l with Figures 3f,h): even for the lowest firing rates, the collective activation of a distal synaptic group leads to an NMDA spike and therefore potentiation. In the following simulation, we aimed to combine results for both clustered and distributed synaptic arrangements across different distances from the soma. We therefore divided our basal dendritic tree in different regions depending on the distance from the soma (Figure 3m). For each distance-range, we distributed synapses either randomly among the respective compartments, or they were clustered on one of the compartments. The neurons were activated using Poisson spike trains with different average rates, and we stored the resulting number of unidirectional and bidirectional connections in each case. The mean of these numbers for the distributed and the clustered simulations are summed per distance-range. We ran an equal amount of simulations for all distances. In our neuron, however, there are substantially less compartments for the smallest and largest distance-ranges compared to intermediate ranges. To have a sense of absolute differences of this number of uni-and bidirectional connections across basal dendrites, we further multiplied this number by the amount of compartments in the respective distance-range (Figure 3n).

Finally, while keeping the synapses clustered on either a proximal or distal compartment, we changed the activation pattern. Instead of generating Poisson spike trains simultaneously in both neurons, we now added a temporal structure: the first neuron will always be activated before the second (Figure 3o). In this case, proximally connected neurons will always become unidirectionally connected by potentiating synapses from the first to the second neuron (pre-post activations) while depression synapses from the second to the first (post-pre activations) (Figure 3p,q). Conversely, distally connected neurons always form bidirectional connections since every presynaptic activation generates an NMDA spike in the postsynaptic neuron (Figure 3p,r).

We conclude that the synaptic distribution highly influences the connectivity between neurons. If synapses are not clustered, our model predicts that proximal connections can be strengthened even for relatively low rates (less than 10 Hz) while distal synapses only allow for LTP when the pre- or postsynaptic neuron fires at higher rates (10 Hz or more). However, if synapses are clustered on the same dendritic compartment, distal synapses lead to potentiation even for the lowest rates and independently of any temporal order of neuronal activation. Therefore, we predict that experimentally observed pairs of neurons connected with distally clustered synapses should express mainly bidirectional connectivity, while pairs of neurons that do not show distal clusters would over-represent unidirectional connections. To test our predictions, we re-analysed published data from Markram et al. [Markram et al., 1997]. In that study, the average location of unidirectional connection is at 89.6*µ*m ± 28.6*µ*m whereas the average location of bidirectional connection is at 106.9*µ*m ± 33.7*µ*m (see details in Methods). This data is in qualitative agreement with our prediction of neurons with clusters.

### 4/ Plateau potentials can modulate plasticity at other synapses

The dendritic NMDA spikes are powerful regenerative events, with long plateau depolarisations lasting up to hundreds of milliseconds [Major et al., 2008, Antic et al., 2010]. As they are more easily evoked at the terminal regions of basal dendrites, they undergo substantial attenuation and cause only subthreshold events at the soma. This long subthreshold plateau reduces the amount of depolarisation required to reach the spiking threshold and therefore enables other weak inputs to reach the threshold more easily.

We investigated how the interaction between plateau potentials and subthreshold inputs affects plasticity. In particular, we explored the interplay between STDP and dLTP by connecting three pools of synapses randomly across the basal dendrites. One extra group of synapses is added at the distal part of a thin branch (Figure 4a). The three pools are activated sequentially, but are too weak to induce potentiation by STDP. During the first half of the simulation, the activation of the first pool (Figure 4, brown) is coincident with the activation of the distal group (Figure 4, red). The weak inputs from the first pool paired with the plateau depolarisation now enable the neuron to elicit somatic spikes, which in turn lead to potentiation of synapses belonging to the first pool. During the second half of the simulation, the distal group is activated together with the second pool (Figure 4, yellow) instead of the first. The synapses belonging to the first pool depress again, while those from the second pool, now coupled with the plateau potential, are potentiated (Figure 4d).

**Figure 4.**
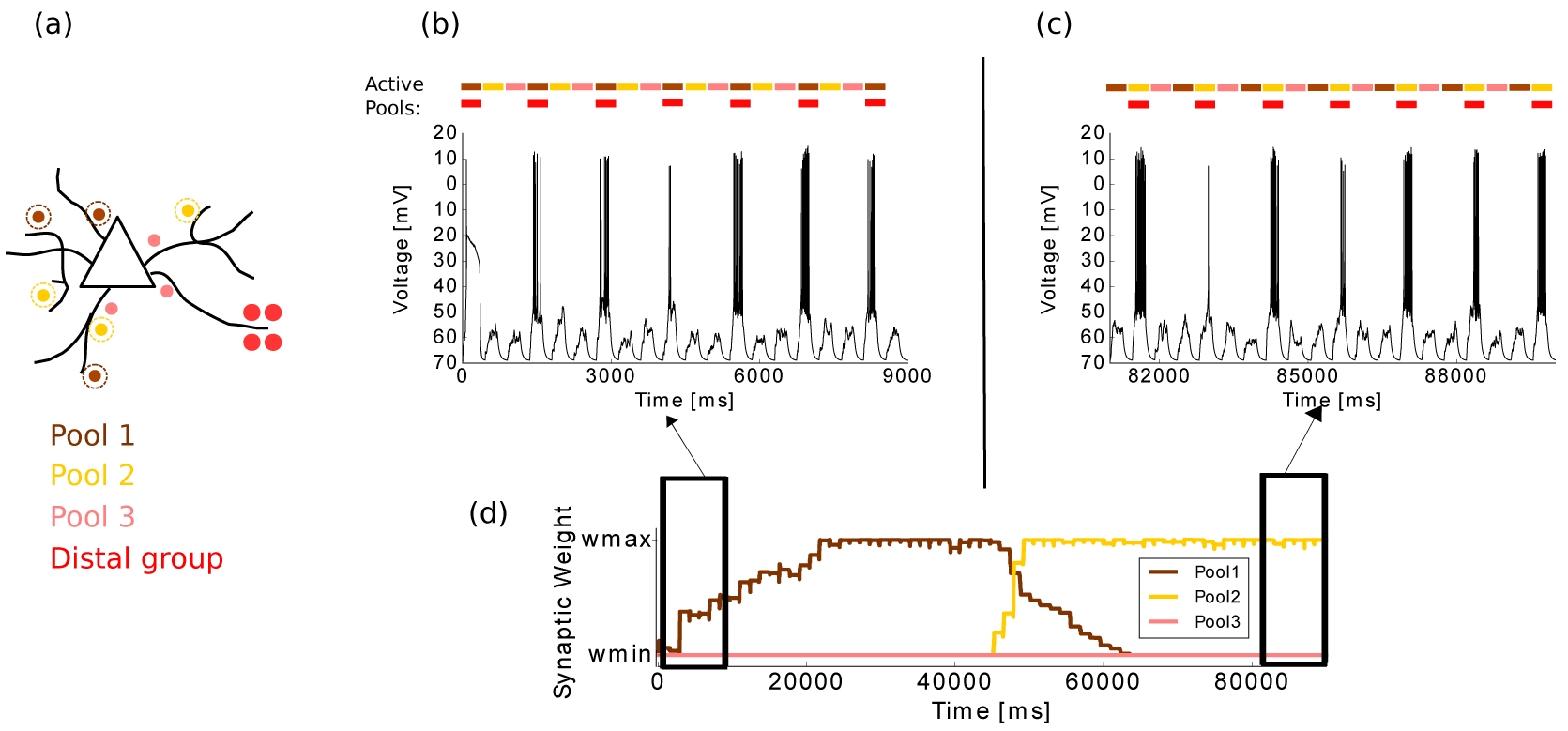
Distally evoked plateau potentials can switch the direction of plasticity at other synapses. (a) Different synaptic groups are connected to the neuron: three pools (brown, yellow, pink) are randomly distributed across the basal dendrites, and one group (red) is connected on one distal compartment. (b) The distal cluster (red) is paired with pool 1 (brown) in the first part of the simulation. The neuron almost exclusively reaches the spiking threshold when these synapses are active together. This leads to potentiation of the brown synapses ((d), first part). (c) The distal cluster (red) is paired with pool 2 (yellow) in the second part of the simulation. The brown synapses depress again and the yellow synapses potentiated ((d), second part). As the distal (red) group evokes an NMDA spike every time it is activated, it remains at the maximum bound.

Our computational model therefore proposes a potential computational role of the these diverse plasticity rules, namely that distal inputs can “gate” or “decide” which proximal inputs will learn.

### 5/ Retention of associations with dLTP in single cells

We then sought to determine whether the dLTP affects the retention of learned associations. In the vSTDP rule, LTD depends on presynaptic activity and postsynaptic depolarisation (see methods). When a group of synapses is activated but fails to evoke a somatic or dendritic spike, the weights will be depressed. This situation can arise when activating only a fraction of associated inputs, leading to depression and in turn unlearning or forgetting of the previously learnt association. However, an association should be robust when presenting only its components, since neurons taking part in multiple assemblies and various sources of noise can lead to an incomplete activation. For example, imagine that we learned the association of coffee, consisting of its features such as color brown, taste, smell, etc. We however don’t forget to associate the color brown with the item coffee even though we see that color much more often than we drink a coffee. As the number of synapses needed for an action potential is larger than those needed for an NMDA spike (Figure 1d), we hypothesize that cooperative synapses allow for an increased retention of associations.

We tested this idea in our detailed model. First, we activated two proximally grouped pools of neurons. We assume that the neuron should respond to the activation of both pools only, but not to the pools separately. For simplicity, the first pool is defined as coding for a circular shape and the second one for a blue color (Figure 6a). We further assume that we encounter many non-blue circular shapes and many non-circular blue objects, while only sometimes observing blue circles. Hence, the separate activation of the circle-pool and the blue-pool will occur much more than the simultaneous activation of both pools (blue circle) (Figure 6b). Even though the simultaneous activations (blue circle) might elicit somatic spikes at the start of the simulation, the multitude of single-pool-activations will depress the involved synapses, ensuring that even simultaneously activating the pools eventually becomes subthreshold (Figure 6b,c). This poses a fundamental limit on retaining cell-assemblies with similar STDP models.

**Figure 6.**
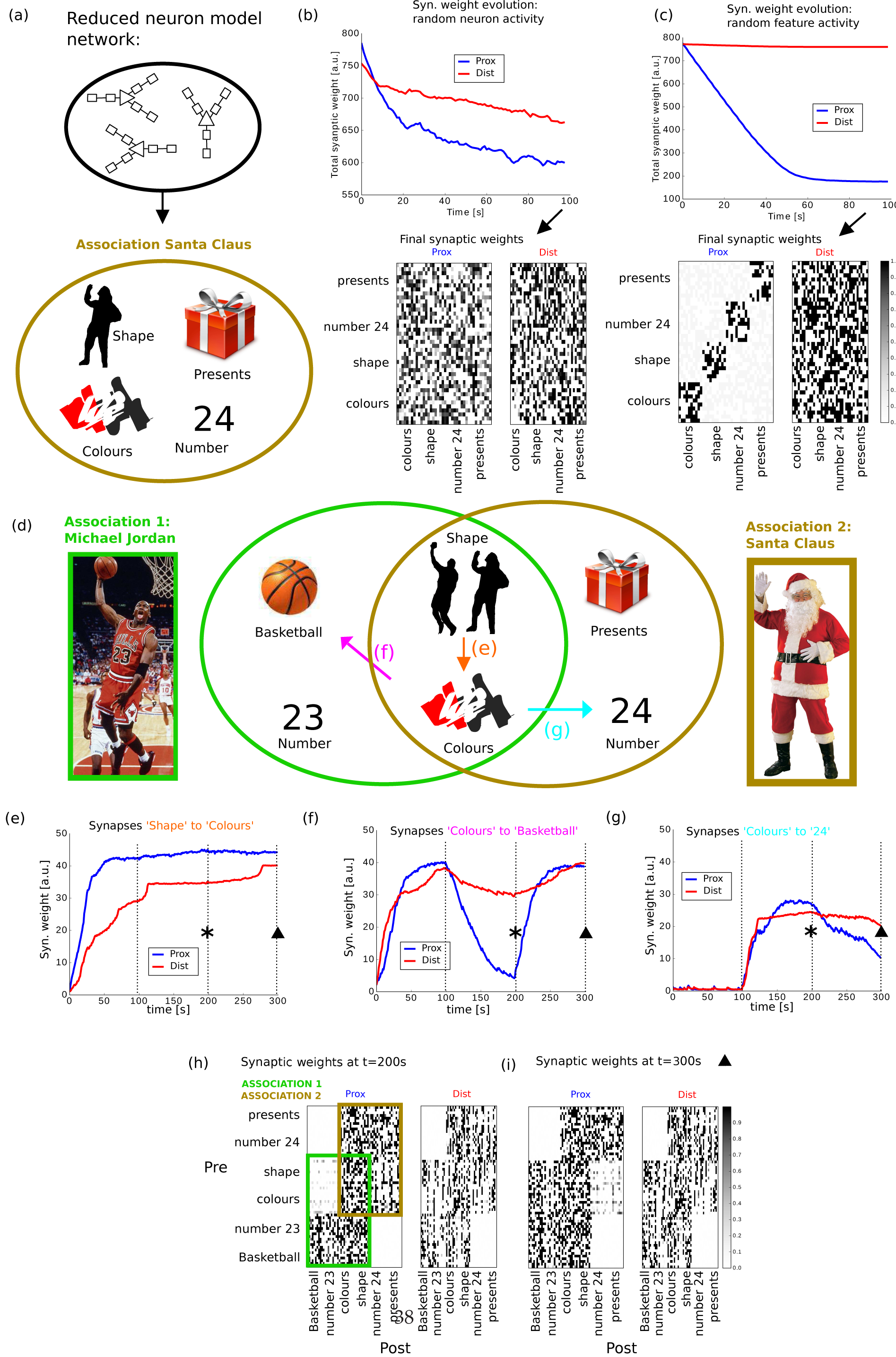
In networks, dLTP can protect previously learnt connections. (a) Top: A neuron model with simplified dendrites is implemented for the network simulations. The model consists of two compartments per dendrite, one representing a proximal compartment and one representing a distal compartment (see also Supplementary). Bottom: A network represents an associative memory (‘Santa Claus’), containing 4 subgroups of neurons which we will call ’features’, in this case they are colours (red, white, black), the shape of a man, presents and the number 24 (Christmas is celebrated on the 24th of December). The weights are all-to-all and initiated at the maximum bound. (b) Top: When randomly choosing 25 percent of the neurons for each activation, both proximal and distal connections are depressing at a similar rate. Bottom: the corresponding final synaptic weight matrix. (c) Top: When randomly choosing a whole feature for each activation, the distal synaptic clusters are always activated together and evoke NMDA spikes. While proximal weights between features depress, distal weights remain close to the initial value. Bottom: corresponding final synaptic weight matrix. (d) A second network consists of two associative memories, ‘Michael Jordan’ and ‘Santa Claus’. Each association is composed of 4 features. The associations share two features (‘Shape of a man’ and ‘Colours red, white, black’), the initial connectivity is all-to-all but at the lower bound. (e,f,g) During the first 100s, the ‘Michael Jordan’ association is activated ten times more frequently than the ‘Santa Claus’ association. Proximal and distal synaptic weights between neurons of the ‘Michael Jordan’ association are strengthened (e,f). From 100s to 200s, we reverse the presentation probability: the ‘Santa Claus’ association is presented ten times more frequently. The synaptic weights between neurons of this association are now maintained or strengthened (e,g). The proximal synapses that are exclusively part of the ‘Michael Jordan’ association are now weakened, while the corresponding distal synapses remain more stable (f). During the final 100s, the same protocol as in the initial 100s is followed, activating mostly ‘Michael Jordan’ association. The proximal synapses that are exclusively part of the ’ coffee’ association are now weakened, while the corresponding distal synapses remain more stable (g). (h,i) Weight matrix after 200s (panel (h) and asterisk in (e,f,g)) and 300s (panel (i) and triangle in (e,f,g)) of the simulation. Notice the difference in the proximal weights (left panels) from features ‘Shape’ and ‘Colours’ to features ‘23’, ‘basketball’, ‘24’ and ‘presents’. The distal weights (right panels) remain similar

When two pools (circular shape and red color, Figure 6d) are located at the distal parts on basal dendrites, each activation of a pool elicits an NMDA spike, therefore keeping the synapses at the maximum weight (Figure 6e,f). Importantly, keeping the weights at full strength does not require any action potentials in this case, and the separate activations of each feature remain subthreshold and do not interfere. This allows the neuron to reliably produce output in the event of a red circle only (Figure 6e). Our example shows how dLTP provides a mechanism to maintain associations between various inputs by protecting the synapses to be overwritten by other activity. Underlying this behaviour is a crucial difference between dLTP and STDP. When using only STDP, the neuronal output also provides the learning signal for potentiation. Indeed, an action potential (output) is necessary for the induction of any weight change (learning). This is in contrast with dLTP, where the potentiation is completely decoupled from the output (action potential generation) but instead relies on NMDA spikes. Therefore, the learning can happen independently from the output, allowing the different features of the learnt association to be activated separately without leading to depression.

### 6/ Memory retention in networks of neurons with dendrites

To investigate the network effects of dLTP, we implemented a reduced neuron model. In this model, each dendrite consists of only two compartments, one mimicking the proximal part of basal dendrites and the other mimicking distal regions (top panel in Figure 6a and see methods and Supplementary Figure S4).

Building further on the results in section 5, we wondered how ongoing activity affects memory retention in networks. In the first network, we implemented 4 groups of neurons, which we will assume to be 4 features that constitute one ‘association’. For example with ‘Santa Claus’, one could associate the features 24 (the date celebrating Christmas eve), colours red/white/black, the shape of a man and presents (bottom panel in Figure 6a). The network neurons are all-to-all connected and the connections are randomly distributed across distal and proximal compartments. Importantly, the distal connections coming from neurons of the same feature are always clustered on the same distal compartment post-synaptically. We then simulated random ongoing activity by choosing on average 25 per cent of the total amount of neurons for every activation event. Even though the total amount of proximal weights drops faster than the distal weights in the beginning of the simulation, eventually both proximal and distal weights depress at similar rates (Figure 6b). The rate of depression is slow, as depression events between two neurons are only about three times more likely then potentiation events: depression occurs when the presynaptic neuron is active while the postsynaptic neuron remains silent (*P*_pre_ · *P*_no post_ = 1/4 · 3/4 = 3/16), potentiation occurs when both presynaptic and postsynaptic neurons are active (*P*_pre_ · *P*_post_ = 1/4 · 1/4 = 1/16). We then repeated this simulation, but instead of choosing individual neurons arbitrarily for each activation, we randomly chose and activated a whole feature. Using the example above, we can imagine to encounter red/white/black colors, number 24, presents and shapes of men at occasions other then when thinking about Santa Claus. Since the neurons from these different features are never activated together, proximal weights between different features are weakened. However, the activated features now always correlate with distally projecting clustered synapses, and NMDA spikes will be evoked more easily. As a result, we find that the distal weights do not depress substantially compared to the proximal weights (Figure 6c).

The ongoing activity as in the previous simulation might be caused by the activation of different associative memories sharing groups of neurons. We therefore explored how the learning and relearning of such associations affect each other. We divide a network into 2 associative memories, for example one ‘Michael Jordan’ and the other ‘Santa Claus’. Each consists of 4 groups of neurons, representing different features of the association. Both Michael Jordan and Santa Claus share colours and shape features while having 2 unshared features each, resulting in a total of 6 features (Figure 6d).

The associative memories have a different probability of being activated, and we are interested in how these ongoing activations interfere with previously learnt connections. We hypothesise that dLTP, which allows a neuron to disconnect its output from its learning, enables the retention of previously learned connections in situations where STDP would not. We initialize our connections at the minimum weights and neurons are all-to-all connected. The connections have a 50 percent probability of being proximal or distal. Furthermore, distal connections arriving from neurons that are part of the same feature, are clustered onto the same distal compartment of postsynaptic neurons.

In a first phase of the simulation, ‘Michael Jordan’ has a tenfold higher probability to be activated then ‘Santa Claus’. In this phase, both distal and proximal connections between neurons activated by ‘Michael Jordan’ are strengthened (Figure 6e,f left side). In the second phase, we reverse the probabilities and ‘Santa Claus’ is now ten times more likely to be activated than ‘Michael Jordan’. Since these memories share half of their neurons, learned connections can be erased. More specifically, only presynaptic activity without postsynaptic activity leads to depression in our model (see methods), and hence the connections from the shared neurons to the unshared neurons of the first association are expected to be erased. However, the distal connections allow NMDA spikes to be evoked post-synaptically, and hence are not depressed (Figure 6f, middle). This leads to a partial retention of the learned associations, where the distal connections protect previously strengthened synapses from being completely erased. In the final phase, we reversed the probabilities again, observing a similar strong depression of proximal weights from the shared neurons to the unshared neurons of the Santa Claus association, while the distal weights remain close to the value at the start of the phase (Figure 6g, right).

Our simulations suggest that dLTP allows a subsection of strengthened weights to be maintained for a longer time compared to STDP. Due to this mechanism, a trace of a previously learned memories could remain present, even when such a memory has not been activated for a long time. The dLTP protects the weights from being weakened by ongoing activity, while synapses unable to evoke dLTP are depressed. The strength of the latter type of synapses therefore depends on a more recent history of activations, favouring associations which are more ‘fresh’ in the memory.

## Discussion

In our work, we investigate how dendrites provide various plasticity mechanisms to a neuron, and how their relative importance depends on the location of the synapse along the dendrite. We modelled two detailed multicompartmental neurons reconstructed by [Acker and Antic, 2009] and [Branco et al., 2010], with active conductances, AMPA and NMDA mediated synapses, and a voltage-dependent plasticity rule [Clopath et al., 2010]. As we currently have no knowledge of experimental data showing otherwise, we assumed the same plasticity rule in both the layer 2/3 and the layer 5 neuron. Our model reproduces the experimentally observed plasticity compartments along single dendrites [Gordon et al., 2006, Weber et al., 2016] and the distance-dependence of STDP [Froemke et al., 2005, Sjöström and Häusser, 2006, Letzkus et al., 2006]. This distance-dependence was shown for low pairing frequencies in [Froemke et al., 2005, Letzkus et al., 2006]. In our model, at low rates this effect seems negligible. The discrepancy could be explained by considering differences in induction protocol: experimentally a spike might be induced by a stronger extracellular shock compared to our simulations. In the experiments, each spike would then be surrounded by a larger depolarisation and since we use a voltage-dependent plasticity rule, this would emphasize the difference in plasticity between proximal and distal locations.

Experiments have shown that proximal regions difficultly allow for NMDA spikes while more distal regions favour dLTP [Golding et al., 2002, Kampa et al., 2006, Gordon et al., 2006, Gambino et al., 2014, Brandalise and Gerber, 2014, Cichon and Gan, 2015]. However, it is hard to quantify the importance of dLTP from this data. Our model allows us to compare the prevalence dLTP and STDP in more detail. Putting together the results of Figure 1 and Figure 2, we firstly see that STDP is present across the whole basal dendritic tree in our model, although potentiation is attenuated distally. Indeed, the innervation by back-propagating action potentials (bAPs) is most powerful in the proximal region on basal dendrites [Nevian et al., 2007, Acker and Antic, 2009]. Secondly, in our model dLTP is possible in more than half of the basal dendritic tree and usually requires less inputs compared to STDP. This mechanism allows for potentiation to occur even when STDP fails, for example when the inputs are too weak to evoke action potentials. These observations would suggest that dLTP and STDP would have comparable importance in basal dendrites. However, another requirement for dLTP is that functionally related synapses are found in spatially proximal locations. It is unclear to what extent such functional synaptic clusters are present on dendrites [Harvey and Svoboda, 2007, Fu et al., 2012, Druckmann et al., 2014, Jia et al., 2010, Chen et al., 2011]. The typical length along which calcium signals spread and dLTP can occur is around 10-20 microns [Schiller et al., 2000, Hill et al., 2013, Weber et al., 2016], which suggests that the synapses can more directly communicate on this scale without the need for action potentials. This form of potentiation would need as few as 2 to 4 synapses within 20 microns of dendritic length [Weber et al., 2016]. Therefore, considering a spine density of about 1-1.5 spines per micron in human cortical basal dendrites [Benavides-Piccione et al., 2013], such clusters do not need to be apparent and neighbouring spines can easily form synapses from distinct signals: different clusters can be intertwined and can mingle with unclustered synapses. In order to arrange synapses with the necessary spatial proximity, structural plasticity might play an additional important role [Fu et al., 2012]. Indeed, similar synaptic arrangements have been achieved in theoretical models by a combination of cooperative and structural plasticity in [Mel, 1991, Poirazi and Mel, 2001].

Homeostatic mechanisms could also provide a strengthening of weak inputs unable to evoke action potentials. It is however important to note that this is only achieved in an unspecific manner: when most inputs to a neuron are removed for a long period, all synapses should be strengthened simultaneously [Turrigiano and Nelson, 2004]. This is in contrast with the highly specific potentiation due to cooperation of nearby spines. While the generation of NMDA spikes is in principle required for this type of potentiation with the current model parameters, a recent study in hippocampus revealed dLTP occurring without dendritic spikes [Weber et al., 2016]. To account for this type of potentiation, we would need to refine our plasticity rule. We point out that the resulting power of dLTP relative to STDP would be further increased, as even less synaptic inputs would be required by the former, and therefore the results would not change qualitatively. This mechanism of plasticity could be additionally gated by brain-derived neurotrophic factor (BDNF), as observed in [Gordon et al., 2006]. Figure 2f uncovers that, in our model, synaptic potentiation and depression are not quite balanced at distal locations. The dLTP is exceptionally powerful due to the long-lasting nature of the NMDA spike. Even though it is likely that some saturation is present, we chose not to modify our model further (except for Figure 6) as we have no experimental constraints to do so at this point. It would be interesting to investigate how LTP depends on the duration of an NMDA spike and how it compares to STDP, allowing us to further refine our plasticity model. In the network simulations (Figure 6) we wanted to show that the observed effects are not due to a disproportionately strong LTP, but rather due to the decoupling of the learning from the neuronal output by dLTP. Therefore, we balanced the distal LTD-LTP strength by reducing the dLTP amplitude by a factor of 5. The synaptic depression and STDP at both proximal and distal compartments remained identical.

Inspired by experimental findings, Mehta [Mehta, 2004] proposed a dendritically constrained learning rule which is independent of somatic spikes and similar to dLTP. The author discussed how his proposed rule should give rise to functionally clustered synapses and wondered whether learning rules could be different in proximal and distal regions of the dendritic tree. However, the proposed learning rule was not implemented. In a later computational study, an NMDA-receptor-dependent plasticity rule at single synapses was used to investigate location dependence of plasticity [Kumar and Mehta, 2011]. In this study, the authors showed that individual synapses are tuned to an optimal input frequency when taking into account saturating effects of calcium-dependent LTP, and that this frequency is higher for more distal synapses. A similar conclusion is reached in our model for unclustered synaptic configurations. More precisely, if experiments show a negligible amount of clusters, we predict that distal synapses need higher rates than proximal synapses to be strengthened. This would result in distal regions of a neuron having higher average input rates compared to proximal regions. Furthermore, our results show a substantially different connectivity arising from clustered synapses compared to unclustered synaptic arrangements. If experiments would show a substantial amount of clustering, we predict that distally clustered synapses can strengthen even at low rates. This would imply an over-representation of bidirectional connections for neurons connected with distal clusters, while proximally connected neurons would express more unidirectional connections (Figure 3). We did not consider combinations where synapses of one neuron are clustered distally while the synapses from the other neuron are not. However, one could imagine taking an intersection between the results of distally clustered synapses (Figure 3i) and the other cases (Figure 3c,d,g,h,k). Such mixed cases would result in an increase of both unidirectional connections and bidirectional connections, depending on which neuron fires at the highest rate. We re-analysed published data from Markram et al. [Markram et al., 1997], which is in qualitative agreement with our prediction of the presence of clustered synapses. It would be interesting to see if future experimental data shows a dependence of input rate and connectivity on the synaptic distance from the soma.

We further showed how the plateau depolarisation in the soma, caused by dendritic spikes, enables other weak inputs to reach the spiking threshold (Figure 4). In this way, the more distally located inputs can determine the direction of the plasticity at other synapses. Such an interplay where the strengthening and weakening of synapses depends on the coincidence with a dendritic spike could be exploited by having the dendritic spike as a teacher signal. Similar mechanisms, where inputs on a region of the dendritic tree can control the plasticity at a distinct area have been reported in hippocampus [Dudman et al., 2007, Takahashi and Magee, 2009, Brandalise and Gerber, 2014] and explored in theoretical models [Urbanczik and Senn, 2014]. Moreover, it was shown that the modulation of gain curves between different regions on a single branch is asymmetric and location dependent [Behabadi et al., 2012], revealing a complex interplay between distal and proximal inputs. We then provided an example of how a neuron can use dLTP to maintain learnt associations, which would otherwise be depressed by an STDP mechanism (Figure 5).

**Figure 5.**
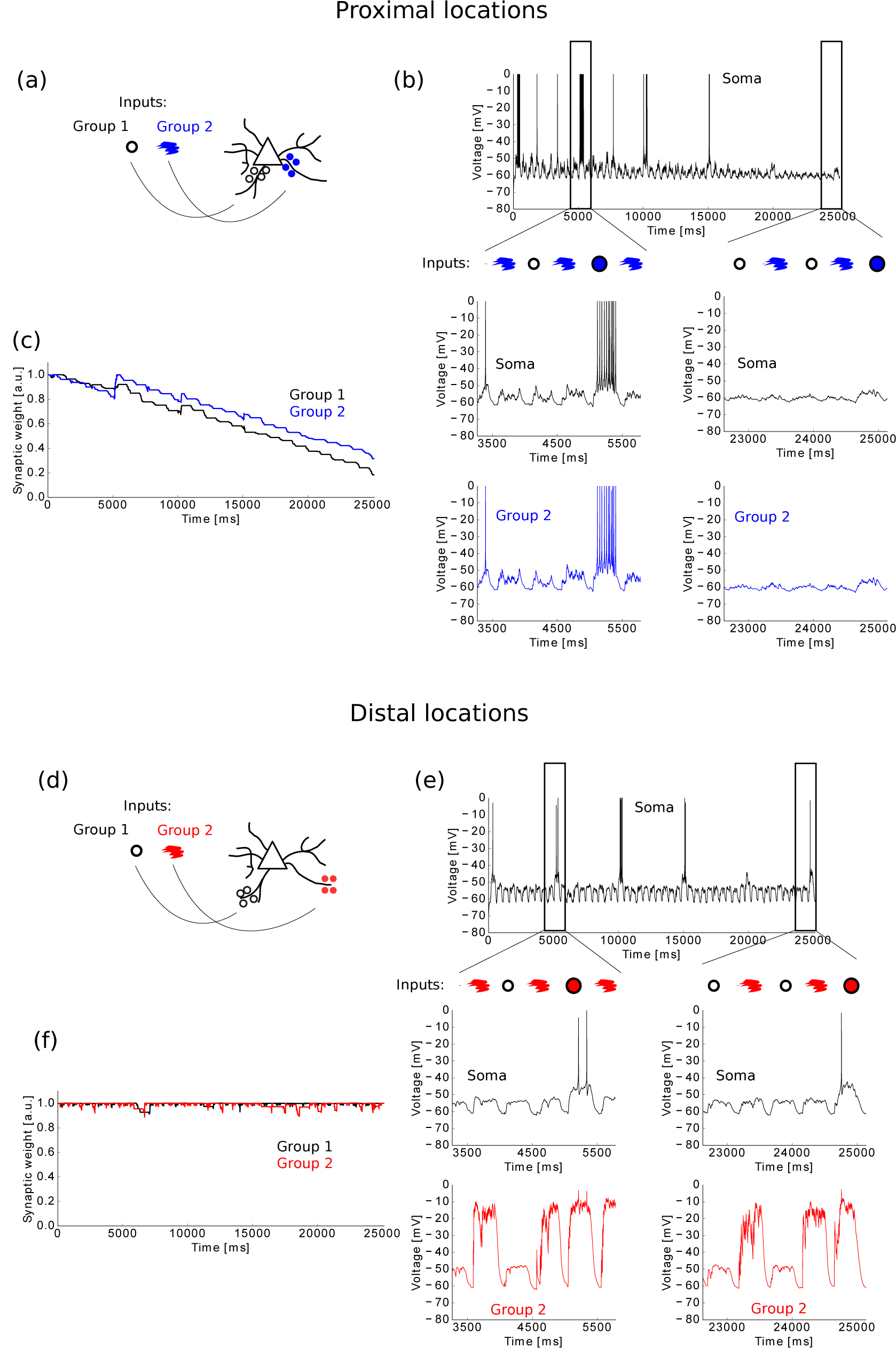
Subthreshold inputs are maintained by dLTP. (a) Cartoon of the simulation: two groups of synapses are placed at proximal compartments. Group 1 codes for a circle, group 2 for the colour blue. While the groups are often activated separately, the neuron should only generate an output in the rare event of a blue circle (i.e. both groups active simultaneously). (b) Top panel: somatic trace during the simulation. Middle panels: detail of the somatic trace during initial half and final half of the simulation. While a blue circle leads to action potentials initially, it eventually becomes subthreshold. Bottom panels: dendritic traces at the proximal location of cluster 2, corresponding to the somatic traces in the middle panel. (c) Evolution of synaptic weights of both groups during the simulation. The individual activations of the groups lead to depression, the simultaneous activations result in potentiation initially, but insufficient to keep the synapses at the maximum weight. Finally, the simultaneous activations become subthreshold and the weights depress towards the lower bound. (d,e,f) Analogous to (a,b,c), but connecting the groups at distal locations on basal dendrites. The individual activations of the groups now lead to NMDA spikes (see panel e, bottom, for the second group), keeping the weights at the upper bound (panel f) and allowing the neuron to maintain its red-circle representation (panel e, top and middle).

Finally, we implemented a reduced neuron model, with two compartments per dendrite. Similar simplified models with reduced dendrites have been used previously, highlighting the advantage of dendrites for computational purposes [Legenstein and Maass, 2011, Behabadi et al., 2012, Jadi et al., 2012, Cazé et al., 2012, Urbanczik and Senn, 2014, Hawkins and Ahmad, 2016, Kastellakis et al., 2016, Thalmeier et al., 2016, Brea et al., 2016]. We showed in networks of reduced neurons how dLTP allows a subgroup of strengthened synapses to remain strong, while they would be weakened under STDP. In this way, dLTP can prolong the retention of associative memories. For these network simulations, we assumed that distal connections arriving from neurons of the same group are already clustered onto the same distal compartment of postsynaptic neurons. It is plausible that the formation of the clusters itself requires an additional mechanism, like structural plasticity, and it would therefore likely take a longer time for strong distal clusters to form. Similar mixtures of both fast and slow learning rates have been discussed in the context of memory retention and the plasticity-stability dilemma [Fusi et al., 2005, Roxin and Fusi, 2013]. It was shown that both a large initial learning due to a fast learning mechanism and a long retention due to a slow mechanism were possible. Our results could be regarded as a new implementation of such a combination of learning rates. Note that when we speak about memory retention in this article, we mean the protection of being overwritten by other activity and not synaptic consolidation [Uwe Frey and Morris, 1997]. However, a distal cluster could have a similar role to synaptic consolidation and could therefore help in the trade-off between plasticity and stability. While we mainly focussed on the prolonged retention, we proposed asynchronous activity and inhibition as mechanisms enabling the depression of distally clustered synapses. These mechanisms could then be exploited to maintain sparse representations, by weakening the distally clustered connections when they should be unlearned.

In this study, we did not consider various other plasticity mechanisms: intrinsic and homeostatic plasticity [Triesch, 2007, Turrigiano, 2011, Turrigiano, 2012, Yger and Gilson, 2015], structural plasticity [Caroni et al., 2012], heterosynaptic plasticity [Chistiakova et al., 2014, Chistiakova et al., 2015] and branch strength plasticity [Losonczy et al., 2008, Makara et al., 2009, Legenstein and Maass, 2011]. Moreover, we only considered basal dendrites, and different learning rules might be present in apical tuft dendrites [Kim et al., 2015, Sandler et al., 2016], especially in cortical layer 5 pyramidal neurons which are known to possess a powerful calcium spike initiation zone in this region [Schiller et al., 1997]. Understanding how this spike interacts with plasticity could provide further insight into the interaction of bottom-up inputs arriving on the basal dendrites and top-down feedback arriving at the tuft. Furthermore, we neglected inhibitory synapses, which are known to target specific regions of the dendritic tree [Markram et al., 2004, Bloss et al., 2016], affect the neuronal output in a location-dependent way [Jadi et al., 2012, Gidon and Segev, 2012] and can influence the back-propagating actionpotential and hence synaptic plasticity [Wilmes et al., 2016].

To conclude, our simulations confirm that in the distal regions on basal dendrites, a small amount of synapses can cooperate to evoke NMDA spikes and trigger LTP. This type of potentiation is independent of action potentials and therefore decouples the learning from the neuronal output. We showed that the connectivity between pairs of neurons is highly influenced by the synaptic arrangement, and made experimentally testable predictions. We further investigated how NMDA spikes can switch the directionality of plasticity at other synapses, and how a longer retention of learned associations arises in networks of neurons with active dendrites, while still allowing the same neurons to form new associations.

## Methods

For all simulations of the biophysical neurons, we used the Brian2 neuron simulator [Goodman, 2008] in Python. The network simulations were developed in Python. The codes will be posted on ModelDB (https://senselab.med.yale.edu/modeldb/) after publication.

## Biophysical neuron models

### Neuron Morphology and passive properties

We used a morphological reconstruction of a Layer 5 and a Layer 2/3 pyramidal neuron of mouse visual cortex, reconstructed by Acker et al. [Acker and Antic, 2009] and Branco et al. [Branco et al., 2010] respectively. The HOC files were converted into an SWC file using the free software NLMorphologyConverter (http://neuronland.org/NLMorphologyConverter/NLMorphologyConverter.html). For this purpose, the original soma of the Layer 2/3 model (21 compartments) was replaced by a single-compartment spherical soma with approximately the same volume (radius of the sphere= 8 µm). The total number of compartments in the Layer 5 and Layer 2/3 model were 1181 and 488 respectively.

The passive parameters for the neuron were:

|  |  |  |
| --- | --- | --- |
| Timestep | $dt$ | 25 $\mu\text{s}$ |
| Leak conductance | $g_L$ | 0.04 $\text{mS cm}^{-2}$ |
| Leak potential | $E_L$ | -60 mV |
| Membrane capacitance | $C_m$ | 1 $\mu\text{F cm}^{-2}$ |
| Axial resistance | $R_a$ | 90 $\Omega \text{ cm}$ |

To account for spines, we increase the leak conductance and membrane capacitance by factor of 1.5 for distances larger than 50 microns, as in [Acker and Antic, 2009]. The value of *E*_*L*_ was chosen to ensure a resting potential of the neuron of -69 mV in the Layer 5 and -72 mV in the Layer 2/3 neuron.

### Ion channels

The active conductances in our model were adapted from [Acker and Antic, 2009], a study in which a fit was made to match experimental data on action potential backpropagation in basal dendrites of layer 5 pyramidal cells. The most prominent differences between layer 5 and layer 2/3 pyramidal neurons are found in the apical dendrites: the Ca^2+^-spike initiation zone and Ih current are almost completely absent in layer 2/3 neurons [Waters et al., 2003, Larkum et al., 2007, Palmer et al., 2014]. These results indicate that the apical tuft dendrites of layer 2/3 neurons might be more similar to the basal dendrites. Nevertheless we chose to distribute synapses solely across the basal dendritic tree in all our simulations.

We further modified the distribution of the channels in the soma and axon from [Acker and Antic, 2009] in order to have a spike initiation in the initial part of the axon instead of the soma. All equations governing the channel dynamics were identical as in [Acker and Antic, 2009], but translated from the HOC files into python with Brian2. As in [Acker and Antic, 2009], most temperature dependencies are removed from the ion conductances, except for the high-threshold calcium channel. The conductance of the latter should therefore be multiplied by a factor of 2.11 for a fair comparison with other conductances, as we used a temperature of 32 degrees Celsius. We point out that in most models with the NEURON simulator, the conductances are temperature dependent, which one should take into account when comparing with the current work. Full details on the ion channel dynamics and distributions can be found in [Acker and Antic, 2009], a summary of the ion channel distributions in our model can be found in the tables below. For simplicity, we will use the following abbreviations:

- Voltage-gated Na^+^ channels : Na_v_
- Delayed-rectifier K^+^ channels: K_v_
- A-type K^+^ channel: Ka_v_
- High-threshold Ca^2+^ channels: Cah_v_
- Low-threshold Ca^2+^ channels: Cal_v_

### Soma

|  |  |
| --- | --- |
| $Na_v$ | $200 \text{ pS } \mu\text{m}^{-2}$ |
| $K_v$ | $400 \text{ pS } \mu\text{m}^{-2}$ |
| $Ka_v$ (proximal type) | $300 \text{ pS } \mu\text{m}^{-2}$ |
| $Ca_h_v$ | $4 \text{ pS } \mu\text{m}^{-2}$ |
| $Ca_l_v$ | $1 \text{ pS } \mu\text{m}^{-2}$ |

**Basal dendrites** Sodium and potassium channel densities linearly depend on the distance d to the soma in microns.

|  |  |
| --- | --- |
| $Na_v$ | $(150 - 0.5*d*\mu\text{m}^{-1}) \text{ pS } \mu\text{m}^{-2}$ |
| $K_v$ | $40 \text{ pS } \mu\text{m}^{-2}$ |
| $Ka_v$ | $(150 + 0.7*d*\mu\text{m}^{-1}) \text{ pS } \mu\text{m}^{-2}$ |
| $Ca_h_v$ | $0.4 \text{ pS } \mu\text{m}^{-2}$ |
| $Ca_l_v$ | $0.1 \text{ pS } \mu\text{m}^{-2}$ |

Apical dendrites

|  |  |
| --- | --- |
| $Na_v$ | $250 \text{ pS } \mu\text{m}^{-2}$ |
| $K_v$ | $40 \text{ pS } \mu\text{m}^{-2}$ |
| $Ka_v$ | $300 \text{ pS } \mu\text{m}^{-2}$ |
| $Ca_h_v$ | $0.4 \text{ pS } \mu\text{m}^{-2}$ |
| $Ca_l_v$ | $0.1 \text{ pS } \mu\text{m}^{-2}$ |

The Ka_v_ channel density in all dendrites is further divided into proximal and distal A-type potassium channels (full details in [Acker and Antic, 2009]).

**Axon** In order to mimic spike initiation in the axon, the initial two segments of the axon have a K_v_ density of 40 and 100 pS µm^−2^ respectively and a Na_v_ density of 8000 and 7000 pS µm^−2^ respectively. The rest of the axon has a constant K_v_ density of 500 pS µm^−2^ and Na_v_ density of 5000 pS µm^−2^. There are no calcium conductances in the axon, except in the first two compartments where they have the same values as in the soma. Additionally, a low-threshold activated, slowly inactivating potassium current is added in the first four compartments of the axon, and is zero otherwise. As this study focuses on plasticity in dendrites, the axonal ion distributions were chosen purely to allow for a spike generation and not for biological plausibility.

### Synapses

Synapses consist of both AMPA and NMDA channels. Except for Supplementary Figure S3, the maximal conductance of NMDA receptors is always double of the AMPA conductance, as was used in [Branco and Häusser, 2011] to reproduce the integration gradient along dendrites. This ratio is within the observed experimental range [Watt et al., 2000]. However, it is usually assumed that only AMPA receptors are influenced by synaptic plasticity and NMDA receptors underly homeostatic scaling. To account for this, we re-ran most simulations with only the AMPA receptors plastic. As we started our simulation with the same AMPA to NMDA ratio, the outcomes remained similar. In Supplementary Figure S3, we give an example of an dendritic spike using a lower amount of NMDA receptors, and show how its duration is reduced. This will in turn reduce the strength of potentiation, without changing our results qualitatively.

An activated synapse will result in an instantaneous rise of both AMPA and NMDA conductances by an amount of *g* = *w* · *g*_max_, where w is the synaptic weight and *g*_max_ is the maximal conductance for either AMPA or NMDA. This is followed by an exponential decay with the respective time-constants for AMPA and NMDA:

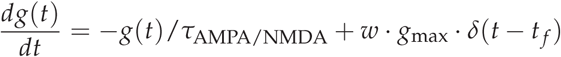

*δ*(*x*) is the Dirac delta function, *t* _*f*_ is the presynaptic firing time and *τ*_AMPA/NMDA_ is the synaptic time constant of AMPA or NMDA receptors. The current flowing through AMPA receptors is modelled by

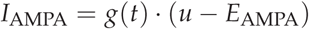

while the NMDA current is given by

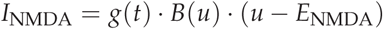

*B*(*v*) describes the voltage-dependence of the magnesium block in NMDA channels [Jahr and Stevens, 1990],

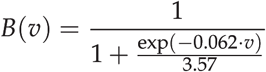

All parameters are below:

|  |  |  |
| --- | --- | --- |
| AMPA time-constant | $\tau_{\text{AMPA}}$ | 2 ms |
| AMPA reversal potential | $E_{\text{AMPA}}$ | 0 mV |
| NMDA time-constant | $\tau_{\text{NMDA}}$ | 50 ms |
| NMDA reversal potential | $E_{\text{NMDA}}$ | 0 mV |
| maximal AMPA conductance | $g_{\text{AMPA}}$ | 1.5 nS |
| maximal NMDA conductance | $g_{\text{NMDA}}$ | 3 nS |

### Plasticity

The plasticity rule used is based on the voltage-based STDP rule from Clopath et al. [Clopath et al., 2010] without homeostasis, where the synaptic weights *w*_*i*_ follow

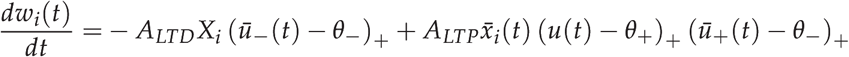

With

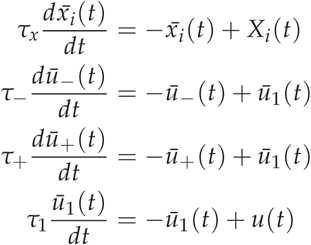

All parameters were in the model were chosen manually in order to reproduce the plasticity experiments in Sjostrom et al., 2001 [Sjöström et al., 2001] at both proximal and distal locations (see Figure 2). We were especially interested in the differences between proximal and distal locations and therefore we aimed for a qualitative agreement with the experimental data. The most important difference with the original model [Clopath et al., 2010] is that the voltage *u*(*t*) used for our implementation was the local voltage at the dendritic compartment, and not the somatic voltage. The voltage *u*(*t*) in each compartment is generated by the Brian2 simulator.

Another difference with the original model in [Clopath et al., 2010] is the extra low-pass filter *ū*_1_, implemented to introduce a small delay in the low-pass filters *ū*_+_ and *ū*_−_. This was achieved in the original model by filtering *u*(*t* − *∊*).

We refer the reader to [Clopath et al., 2010] for full details and give a short summary here: this rule has a depression term and a potentiation term. The term responsible for depression incorporates a presynaptic spike train *X*_*i*_ and the low-pass filtered postsynaptic depolarisation (*ū*_−_ − *θ*_−_)_+_. The (·)_+_ stands for rectification, i.e. the term is only non-zero if the number within the brackets is positive. *ū*_−_ is a low-pass filtered version of the membrane potential with time constant *τ*_−_, and *θ*_−_ is a threshold for synaptic plasticity. To summarize, this term states that a presynaptic spike will lead to depression if the postsynaptic membrane is depolarised, for example because the postsynaptic neuron fired a spike in the recent past. An isolated post-pre pairing will therefore lead to depression. The term responsible for potentiation includes a low-pass filtered presynaptic spike train 
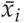
, a low-pass filtered postsynaptic depolarisation (*ū*_+_ − *θ*_−_)_+_ and an instantaneous depolarisation above a potentiation threshold (*u* − *θ*_+_)_+_. The time constant for the low-pass filtered version of the membrane potential *ū*_+_ is given by *τ*_+_. The potentiation threshold *θ*_+_ is high enough to be selective for back-propagating action potentials or NMDA spikes. Potentiation can therefore occur when a postsynaptic spike happens after a presynaptic neuron fired and while the postsynaptic membrane is depolarised, for example because the postsynaptic neuron fired another spike in the recent past as well. Potentiation is therefore possible when there are triplets of spikes (pre-post-post or post-pre-post), which can occur as isolated triplets or during a sustained high rate firing. Secondly, potentiation can occur when an NMDA spike (also called dendritic plateau potential) is generated. These plateau potentials provide a long and sufficiently high depolarisation, leading to potentiation without generating postsynaptic action potentials. The above equations ensure all-to-all plasticity. The parameters are fitted to reproduce the frequency dependence of plasticity, as previously done by [Clopath et al., 2010]:

|  |  |  |
| --- | --- | --- |
| lowpass filter timeconstant 1 | $\tau_1$ | 5 ms |
| lowpass filter timeconstant 2 | $\tau_-$ | 15 ms |
| lowpass filter timeconstant 3 | $\tau_+$ | 45 ms |
| Depolarization threshold for plasticity | $\theta_-$ | -72 mV (L2/3), -69 mV (L5) |
| Depolarization threshold for potentiation | $\theta_+$ | -15 mV |
| Spike trace reset value | $x_{\text{reset}}$ | 5 |
| Spike trace timeconstant | $\tau_x$ | 20 ms |
| Depression amplitude | $A_{\text{LTD}}$ | $4\text{e-}4 \text{ mV}^{-1}$ |
| Potentiation amplitude | $A_{\text{LTP}}$ | $14\text{e-}4 \text{ mV}^{-2}$ |

In all simulations, the minimum and maximum weight for the synapses were 0.01 and 1 respectively, unless stated otherwise. The choice of *θ*_+_, the threshold for potentiation, was made by comparing data from [Sjöström and Häusser, 2006] with [Acker and Antic, 2009]. In the former, it is shown how the attenuation of the bAP results in a transition from potentiation to depression at a distance of about 200 *µ*m, while in the latter it is shown that the bAP amplitude at this distance is roughly 60 mV above the resting potential (−69 or −72 mV in for the layer 5 and layer 2/3 neurons respectively).

We want to point out that all parameters and equations used in our models are the result of fitting procedures or aimed to qualitatively reproduce experimental observations. In doing so, simplifications are necessarily introduced and therefore we do not claim that any value represents a ‘real’ biologically correct value. Instead, the parameters should be interpreted as a set of abstractions that enable a qualitative reproduction of biological phenomena relevant for the scope of this article.

### Reduced multicompartmental model

The reduced compartmental model was implemented in Python. For this purpose, a somatic compartment was coupled to two compartments per dendrite: one representing the proximal section and one the distal section.

The compartments are coupled through there voltage difference and parameters representing axial resistance (details below). The parameters were chosen as to reproduce the L5 detailed model phenomenologically, and might not be the only set of parameters satisfying this behavior. A comparison between both models can be found in Supplementary Figure S4.

Each compartment is modelled using an integrate-and-fire model, based on [Clopath et al., 2010] and where the membrane potential *u*(*t*) of the compartment follows from

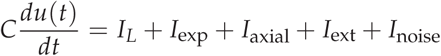

With the following currents:

|  |  |
| --- | --- |
| Leak current | $I_L = -g_L (u(t) - E_L)$ |
| Exponential current | $I_{\text{exp}} = g_L \Delta_T \exp\left(\frac{u(t) - V_T(t)}{\Delta_T}\right)$ |
| Axial currents | $I_{\text{axial}} = -\sum_i g_i \cdot (u(t) - u_i(t))$ |
| External current | $I_{\text{ext}}$ |
| Noisy current | $I_{\text{noise}}(t) = \sqrt{2\tau_{\text{noise}}/\text{ms}} \cdot s(t) \cdot \sigma_{\text{noise}} + \mu_{\text{noise}}$ |
| Inhibitory current | $I_{\text{inhib}}(t)$, see below |

where C is the membrane capacitance, *g*_*L*_ is the leak conductance and *E*_*L*_ is the resting potential. The exponential current models the fast sodium activation in the soma and is not present in the dendritic compartments. Δ_*T*_ is the slope factor for the exponential term describing the sodium current activation, *V*_*T*_ is an adaptive threshold potential which starts at *V*_*T*__max_ after a spike and decays to *V*_*T*__rest_:

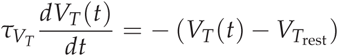

The current entering from or leaving to neighbouring compartments is given by *I*_axial_. The voltage difference between the compartment and each neighbour is multiplied by an appropriate conductance, chosen to reproduce the interaction between soma,proximal and distal compartments of the detailed model. The values for the conductances between soma-proximal and proximal-distal compartments is given below. In order to mimic the difference in efficiency between forward and backward propagation of depolarisations, the conductances are different for both directions:

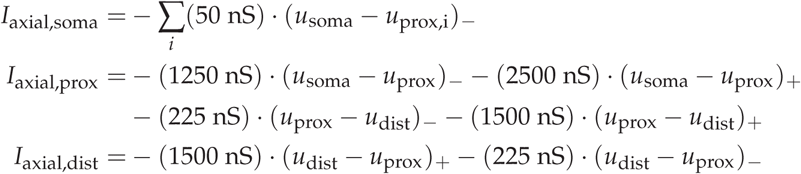

Where we dropped the dependence on t for clarity and (·)_+_ is only nonzero if the value within the brackets is positive, while (·)_−_ is only nonzero if the value within the brackets is negative.

The current *I*_noise_ is coloured noise with standard deviation *σ*_noise_ and mean *µ*_noise_. *τ*_noise_ is the time-constant for the low-pass filtering of the noise and *s*(*t*) is the low-pass filtered version of a Gaussian white noise *ξ* (*t*) with zero mean and unit standard deviation:

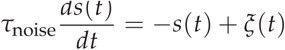

The spike heights at proximal and distal compartments were chosen by taking the mean of spike heights in the compartments of the detailed model shown in Figure 1a. When the voltage in the somatic compartment reaches the spiking threshold (20 mV), a spike is simulated by holding the somatic voltage at 30mV during 1ms. After the spike, the somatic voltage is reset to the reset value *V*_reset_. For every somatic spike, the backpropagation into the dendrites is modelled by holding the dendritic potential at 10mV during 1ms in the proximal compartments, and at −3mV during 1ms in the distal compartments, and with an additional delay compared to the somatic spike of 0.3ms. All other parameters can be found in **Table 1**.

### Synapses

Similar to the detailed model, synapses consist of both AMPA and NMDA channels, where the maximal conductance of NMDA receptors is always double of the AMPAr conductance. An activated synapse will result in an instantaneous rise of the AMPA and NMDA conductances by an amount of *g* = *w* · *g*_max_, where w is the synaptic weight and *g*_max_ is the maximal conductance for either AMPA or NMDA. This is followed by an exponential decay with the respective time-constants for AMPA and NMDA. All parameters are in **Table 1**. We mimic the increased input resistance at distal synapses by multiplying the AMPA and NMDA currents at these compartments by a factor of 2.5.

**TABLE 1:** REDUCED NEURON PARAMETERS

|  |  |  |
| --- | --- | --- |
| Timestep | $dt$ | $250 \mu s$ |
| Leak conductance | $g_L$ | $40 \text{ nS}$ |
| Resting potential | $E_L$ | $-69 \text{ mV}$ |
| Membrane capacitance | $C_m$ | $281 \text{ pF}$ |
| AMPA timeconstant | $\tau_{\text{AMPA}}$ | $2 \text{ ms}$ |
| AMPA reversal potential | $E_{\text{AMPA}}$ | $0 \text{ mV}$ |
| NMDA timeconstant | $\tau_{\text{NMDA}}$ | $50 \text{ ms}$ |
| NMDA reversal potential | $E_{\text{NMDA}}$ | $0 \text{ mV}$ |
| Maximal AMPA conductance | $g_{\text{AMPA}}$ | $60 \text{ nS}$ |
| Maximal NMDA conductance | $g_{\text{NMDA}}$ | $120 \text{ nS}$ |
| Spiking threshold | $\theta$ | $20 \text{ mV}$ |
| Slope factor | $\Delta_T$ | $2 \text{ mV}$ |
| Adaptive threshold rest value | $V_{T\text{rest}}$ | $-50.4 \text{ mV}$ |
| Adaptive threshold max value | $V_{T\text{max}}$ | $-30.4 \text{ mV}$ |
| Adaptive threshold time constant | $V_{T\text{tau}}$ | $50 \text{ ms}$ |
| Spike width | $/$ | $1 \text{ ms}$ |

### Plasticity

The plasticity rule in the reduced model is exactly as in the detailed model, with the only difference that 
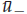
 and
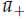
 are defined in the following way:

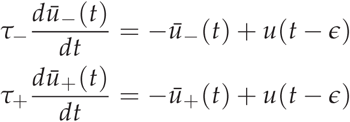

Hence, a small delay is introduced differently in the models: in the reduced model, *∊* is similar to the width of an action potential, we take here 1ms. In the full model, a delay is introduced by an extra low-pass filter *ū*_1_ which is absent in the reduced model. Furthermore, we reduce the amplitude of for dLTP by a factor of 5, in order to have a better balance between depression and potentiation.

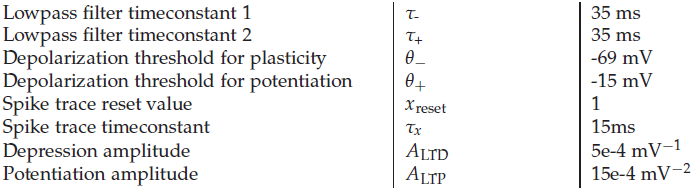

|  |  |  |
| --- | --- | --- |
| Lowpass filter timeconstant 1 | $\tau_-$ | 35 ms |
| Lowpass filter timeconstant 2 | $\tau_+$ | 35 ms |
| Depolarization threshold for plasticity | $\theta_-$ | -69 mV |
| Depolarization threshold for potentiation | $\theta_+$ | -15 mV |
| Spike trace reset value | $x_{\text{reset}}$ | 1 |
| Spike trace timeconstant | $\tau_x$ | 15ms |
| Depression amplitude | $A_{\text{LTD}}$ | $5e-4 \text{ mV}^{-1}$ |
| Potentiation amplitude | $A_{\text{LTP}}$ | $15e-4 \text{ mV}^{-2}$ |

In our network, the value for *A*_LTP_ was reduced by a factor of 5 for dLTP. This was achieved by storing the local voltage value of each compartment during 1.3 ms. We then compared whether the actual value of the local voltage and the saved value 1.3 ms earlier were both above the potentiation threshold *θ*_+_. Since the spike-width was 1ms in our model, this situation could only be caused by an NMDA spike and in that case the potentiation amplitude was reduced. In principle, one could imagine firing rates above 500Hz satisfying the same condition, however the neurons in our network fired at much lower rates.

### Inhibition

To allow higher rates without leading to unbounded activity in our recurrent networks, we approximated an inhibitory current at the soma in the following way. For each neuron, the spikes of all its presynaptic neighbours are filtered with a timeconstant *τ*_inhib_ of 60ms:

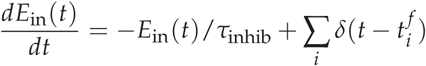

Where 
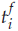
 are the firing times of presynaptic neurons. This filtered trace of incoming excitatory activity *E*_in_ is transformed as follows:

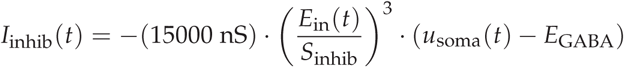

where *I*_inhib_ is the inhibitory current arriving at the somatic compartment, *S*_inhib_ is a scaling constant (150 for Figures 6b-d and 250 for Figures 6e-j), *u*_soma_ is the somatic voltage and *E*_GABA_=-75mV is the reversal potential for GABA.

### Analysis of connectivity in existing data from [Markram et al., 1997]

We reanalysed the data reported in Table 3 in [Markram et al., 1997]. We only used basal and oblique dendrite data, since the plasticity at distal tuft dendrites might be more dependent on calcium spikes [Schiller et al., 1997].

The values for the distance from the soma for unidirectionally connected neurons are 14.7*µ*m, 74.4*µ*m (7x), 75.8*µ*m (14x), 117.7*µ*m, 91.3*µ*m (4x), 134*µ*m (6x), 162.6*µ*m Resulting in an average of (± one standard deviation): 89.6*µ*m ± 28.6*µ*m

The values for the distance from the soma for bidirectionally connected neurons are 73.3*µ*m, 75.6*µ*m (10x), 86.9*µ*m (23x), 97*µ*m (10x), 184.6*µ*m (3x), 157*µ*m (10x), 141.8*µ*m (5x) Resulting in an average of (± one standard deviation): 106.9*µ*m ± 33.7*µ*m

## Simulations

### Figure 1

#### Figure 1a

17 distal and 17 proximal compartments in the neuron model are identified. The distal compartments are further split into two groups: one group consists of distal compartments eliciting NMDA spikes when activating synaptic input as described for Figure 1b below, and the other group consists of compartments where no NMDA spike is elicited with such input. We considered an NMDA spike to be evoked if the local depolarisation crossed a threshold at -15 mV. Using this criterion, 16 out of the 17 distal compartments where classified in the first group and marked with red in the figure. All 17 proximal compartments are shown in blue.

#### Figure 1b

At all compartments shown in Figure 1a, a synapse at the maximum weight *w*_max_ = 1 was connected. For each compartment, the following simulation was performed: the connected synapse was activated 10 times with an interspike interval of 1ms while storing the membrane potential at the soma. This protocol was repeated for interspike intervals of 4ms and 8ms. Figure 1b shows the mean (solid line) and standard deviation (shaded region) of the somatic potential calculated for all proximal compartments (blue) and distal compartments (red).

#### Figure 1c,d

For each compartment of the basal dendrites in our morphological neuron, a number of synapses with fixed weight w=1.5 were activated with interspike intervals of 1ms and the local membrane voltage was recorded during the subsequent 50ms. The simulation was repeated with an increased number of synapses until a threshold at -15mV was crossed. By comparing the local voltage with the somatic voltage, it was determined whether this threshold was crossed due to an NMDA spike or a somatic spike. The number of synapses needed to reach this threshold was stored.

### Figure 2

#### Figure 2b

At the same compartments as shown in Figure 1a, a synapse is activated reproducing the experimental protocol in [Sjöström et al., 2001]. Pairs of pre-post or post-pre spikes separated by 10ms are repeated a further five times at different frequencies of 5Hz up to 50Hz with steps of 5Hz, and for 1Hz. The weight change is multiplied by a factor of 15 to mimic the experimental protocol in [Sjöström et al., 2001]. The initial synaptic weights are 0.5 and the weight change after the six pairings are plotted in function of the frequency. The solid lines represent the means of the simulations in all proximal compartments (blue) or distal compartments (red). The shaded areas represent one standard deviation.

#### Figure 2c

The same pairing protocol as in Figure 2a at 20Hz was repeated for different interspike intervals (1,2.5,5,7.5,10,12.5,15.17.5 and 20ms).

#### Figure 2e

The synapse in one distal compartment is activated 10 times with an interspike interval of 0.1ms and weight w=1.

#### Figure 2f

The compartments shown in Figure 1a are each connected with 10 synapses, all with an initial weight of 0.5. The 10 synapses are activated using a Poisson process with the same average rate, lasting 200ms. This protocol is repeated with various rates between 1Hz to 70Hz.

### Figure 3

We implemented two layer 5 biophysical neurons (using the same morphology) for all simulations. All synaptic weights are initiated at 0.5, and the maximum conductances were reduced to *g*_AMPA_ = 1*nS* and *g*_NMDA_ = 2*nS*. A potentiated synapse was defined as a synapse that had a larger weight value at the end of the simulation compared to the initial value. Bidirectional connected neurons were defined as neurons where the mean of synaptic weights to both neurons was larger at the end of the simulation compared to the start. Unidirectional connected neurons were defined as neurons where only the mean of the synaptic weights to one neuron was larger at the end compared to the start, while the mean of the weights to the other neuron was smaller at the end compared to the start. Unconnected neurons were defined as neurons where the mean to both neurons was smaller compared to the initial value. We realise that looking at the mean value of the weights to discriminate between unidirectional and bidirectional is not completely accurate (one can imagine strengthening of some weights and weakening of others in some situations), but it gives us an adequate way of comparing the synaptic weight changes for the different protocols.

#### Figure 3a-d

Each neuron made 20 synaptic connections on its partner. The synaptic locations were drawn randomly across the whole basal tree. In both neurons, spikes were induced following a Poisson point process with a fixed average rate, and the neurons were simulated for 15s. We then repeated the simulation for all combinations of rates between 0Hz and 20Hz, with 2Hz steps, and stored all the final synaptic weights for every case. We then repeated the whole protocol 20 times with different randomly drawn synaptic locations, which we also stored.

(b) For each of the 20 simulated pairs of neurons, we calculated the mean synaptic distance from the soma and the amount of potentiated synapses present across all combinations of firing rates. (c,d) Two examples of simulation outcomes for different random synaptic locations. See above for our definition of unidirectional and bidirectional connections.

#### Figure 3e-h

Each neuron made 10 synaptic connections on its partner. The synaptic locations were drawn randomly across either the proximal or distal compartments as shown in Figure 1a. In both neurons, spikes were induced following a Poisson point process with a fixed average rate, and we simulated the neurons for 15s. We then repeated the simulation for all combinations of rates between 0Hz and 20Hz, with 2Hz steps, and stored all the final synaptic weights for every case. We then repeated the whole protocol 5 times with different randomly drawn synaptic locations, which we also stored.

(f) For each of the 5 simulated pairs of neurons, we counted the amount of unidirectional or bidirectional connections (see above for how we define these).

(g,h) Two examples of simulation outcomes for different random synaptic locations across proximal compartments (g) or distal compartments (h). See above for our definition of unidirectional and bidirectional connections.

#### Figure 3i-l

Exactly the same as 3e-h, but with the following difference: only one synaptic locations was drawn randomly from either the proximal or distal compartments as shown in Figure 1a, and all 10 synapses were clustered on that compartment.

#### Figure 3m-n

All basal compartments are classified into groups according to the distance from the soma (with increments of 25 microns, except for the final group which comprises of 33 microns). For each distance-range, two neurons are interconnected by 10 synapses each. The synapses are either randomly distributed over compartments in the respective range, or clustered onto one compartment. In both neurons, spikes were induced following a Poisson point process with a fixed average rate, and we simulated the neurons for 15s. We then repeated the simulation for all combinations of rates between 0Hz and 20Hz, with 2Hz steps, and stored all the final synaptic weights for every case. We repeated the whole protocol 3 times (both for the clustered and distributed case) with different randomly drawn synaptic locations. We counted the amount of unidirectional or bidirectional connections (see above for how we define these). To account for the different amount of compartments at different distances from the soma, we multiplied the resulting numbers in each distance-range by the respective number of compartments.

#### Figure 3o-r

Each neuron made 10 synaptic connections on its partner. Only one synaptic locations was drawn randomly from either the proximal or distal compartments as shown in Figure 1a, and all 10 synapses were clustered on that compartment. To simulate temporal order in the activity, we induced spikes only during 10ms in the first neuron, followed by spikes in the subsequent 10ms in the second neuron. These 20ms were then followed by 250ms without any activity, and the whole 270ms were repeated 10 times. The induction of spikes in the first and second neuron during the first 10ms and the second 10ms respectively, were achieved by simulating a Poisson process with average rate of 150 Hz. We point out that although this seemingly high firing rate was only sustained for 10ms (thus on average 1.5 spikes during these 10ms), and over the whole simulation each neuron had an average rate of 5.5 Hz. The whole protocol described above was repeated 5 times with different randomly drawn locations.

(p) For each of the 5 simulated pairs of neurons, we counted the amount of unidirectional or bidirectional connections (see above for how we define these). Note that we only repeated this simulation for one activation rate as opposed to the 55 combinations in the previous simulations, and the uni- or bidirectional connections was always one for proximal or distal synapses respectively.

(q,r) The evolution of mean synaptic weights for one example pair of neurons, in the case where the clusters are proximal (q) or distal (r).

### Figure 4

Three input pools consisting of 10 randomly distributed synapses each are initiated at the minimum weight. Five extra synapses are clustered at a distal compartment on a basal dendrite. These synapses are initiated at the maximum weight and will be paired with one of the other input pools. The activation of the pools or the distal cluster comprises of activating the relevant synapses during 250ms at an average rate of 40Hz. Subsequent activations of pools were separated by a 200ms interval.

During the first half of the simulation (0s-45s), the activations of the first pool are paired with the activation of the distal cluster. During the second half of the simulation (45s-90s), the second pool is paired with the cluster. In order to observe the depression when a pool is not paired with the cluster, the maximum weight is reduced to 0.65 instead of 1. The minimum weight is increased from 0.01 to 0.55, enabling the weakest synapses to still reach the spiking threshold when paired with the cluster. To compensate for the reduced range between the minimum and maximum weight, the amplitudes *A*_*LTD*_ and *A*_*LTP*_ for depression and potentiation are one fifth of the usual values described above.

### Figure 5

#### Figure 5a-c

Two pools of 4 neurons are connected to proximal compartments on basal dendrites (synapses of the same pool are clustered onto the same compartment). The initial weights are at the maximum hard bound 1, and an additional coloured noisy current is injected in the soma:

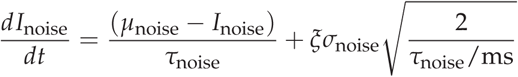

with standard deviation *σ*_noise_ = 10pA, mean *µ*_noise_ = 115pA and *τ*_noise_ = 20ms is the time-constant for the low-pass filtering of the Gaussian white noise with zero mean and unit standard deviation *ξ* (*t*). We alternate the activation of the two pools, with an activation event comprising of Poisson-distributed spikes at an average rate of 30Hz at the member synapses during 350ms. Two subsequent activations are separated by a 150ms interval. Every 10 activations (i.e. 5 for each pool), both pools are activated simultaneously.

#### Figure 5d-f

Analogous as Figures 5a-c, but the two pools are now connected to distal compartments on the basal dendrites. Note that we chose the two distal compartments that most reliably induced somatic spikes when simultaneously activated. Many other combinations failed to do so, but our main result on memory retention of the distally clustered synapses remained in all cases.

### Figure 6

For all simulations of Figure 6, each neuron had 20 dendrites.

#### Figure 6b-d

We implemented 40 neurons, which we divided into 4 groups of 10 neurons. The neurons are all-to-all connected, with weights initiated at the maximum bound for each simulation, and the connections are randomly distributed across distal and proximal compartments. The distal connections coming from neurons of the same feature are always clustered on the same distal compartment postsynaptically. In all neurons, a coloured noisy current was injected in the soma to induce more variability (mean=400pA, std=50pA). Moreover, each neuron had 50 extra non-plastic synapses that were used as inputs.

(c) We chose neurons with a probability of 0.25, and we activated the input synapses to the selected neurons at an average rate of 50 Hz during 50 ms, followed by a 250 ms interval without inputs. The simulation lasted for 100s and the total synaptic strength of the network was stored every second.

(d) We activated the input synapses of one of the groups of 10 neurons (the group was selected randomly) by activating the input synapses of the relevant neurons at an average rate of 50 Hz during 50 ms, followed by a 250 ms interval without inputs. We repeated this during 100s, and randomly chose one of the groups for every activation. The total synaptic strength of the network was stored every second.

#### Figure e-j

We divide a network of 60 neurons into 2 groups of 40 neurons each (‘associative memories’), each group sharing 20 neurons. Moreover, the groups were divided into 4 subgroups of 10 neurons (‘features’), and two of these subgroups were shared. The neurons are all-to-all connected, with weights initiated at the minimum bound for each simulation, and the connections were randomly distributed across distal and proximal compartments. The distal connections coming from neurons of the same feature were always clustered on the same distal compartment postsynaptically. In all neurons, a coloured noisy current was injected in the soma to induce more variability (mean=200pA, std=50pA). Moreover, each neuron had 50 extra non-plastic synapses that were used as inputs.

An activation event consisted of selecting one associative memory and activating the input synapses to the 40 relevant neurons at an average rate of 50 Hz during 50 ms, followed by a 250 ms interval without inputs. The selection of the associative memory was done randomly, but the probabilities changed during the simulation: during the first 100s, the first and second associative memories had 90% and 9% chance of activation respectively, while between 100s and 200s the probabilities were reversed ( 9% and 90% respectively). Finally, between 200s and 300s the activation probabilities were like in the first phase (90% and 9% respectively). The total synaptic strength from subgroups 4 to subroups 1 and 6, and from subgroup 3 to 4 were stored every second. Moreover, the total synaptic weight matrix was stored after 200s and 300s.

## Supplementary Figures

### Supplementary Figures 1 and 2

We refer to the methods for Figure 1 and 2, with the only difference that the analysis is now performed in a layer 2/3 morphology.

### Supplementary Figure 3

We repeated the protocols as in Figure 2e,f, but now with only half the value of the original NMDA conductance.

### Supplementary Figure 4

(c,d) The same protocol as in Figure 2e was repeated for both the reduced and the detailed model. The voltage traces for the compartments shown in (a,b) are stored.

(e,f) A noisy current is injected in the soma of the reduced model and the detailed model. For the reduced model, we used a mean of 200pA and a standard deviation of 2500pA. For the detailed model, we used a mean of 100pA and a standard deviation of 300pA. We remind the reader that these values, especially in the reduced model, are abstract parameters and do not necessarily represent realistic values, but were chosen in order to reproduce a certain behaviour.

(g,h) A stepcurrent of 3000 pA was injected in the soma of our redced model during 3ms. A stepcurrent of 1500pA was injected during 3ms in the soma of our full model. No other stimulation was used and the membrane voltage in the soma, proximal (blue) and distal (red) compartments as shown in (a,b) were stored.

(i) The same protocol as in Figure 2b was reproduced in our reduced model. Figure 2b is shown as panel (j) for reference.

## Author Contributions

J. B. and C. C. planned the research and wrote the paper, J. B. performed the numerical computations.

## Acknowledgements

We thank Arnd Roth for pointing us to the connectivity data of Markram et al. [Markram et al., 1997]. We like to thank Romain Cazé for providing a script for visualising the neuron morphology and Wilten Nicola for comments on an earlier version of the manuscript.

## Supplementary Figures

**Figure S1.**
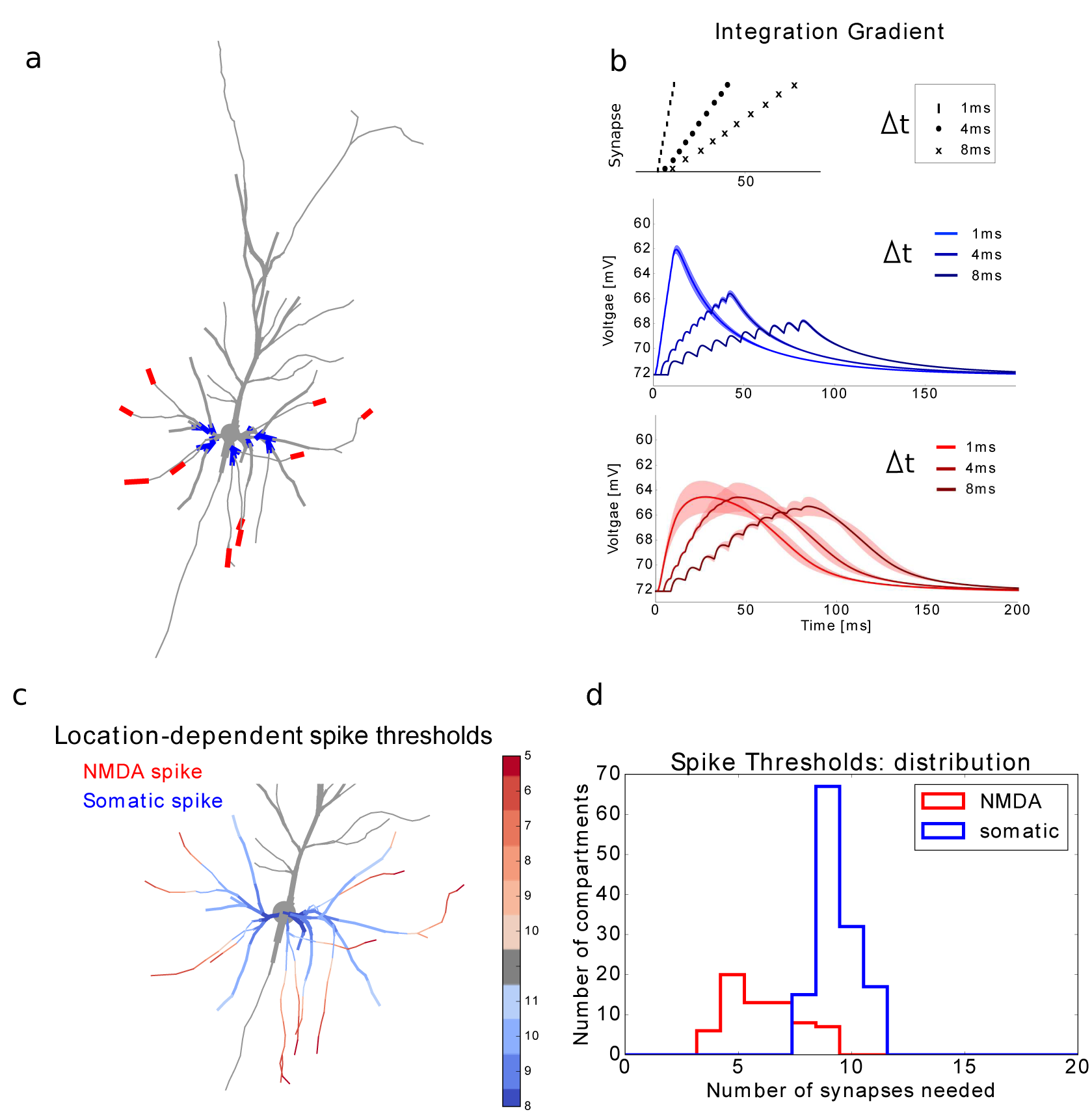
Layer 2/3 pyramidal neuron: location dependence of local and global regenerative events. (See Figure 1, but now for a layer 2/3 neuron morphology)

**Figure S2.**
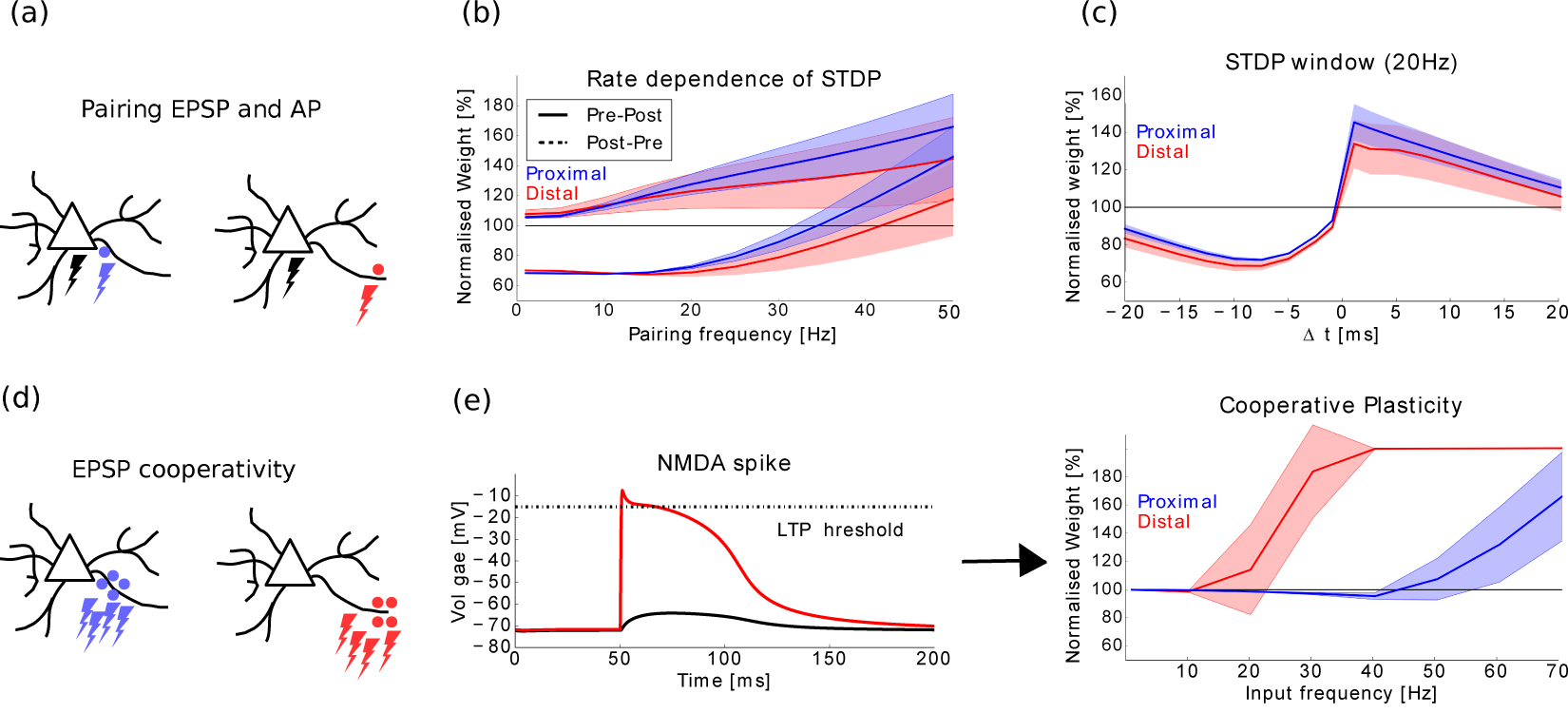
Layer 2/3 pyramidal neuron: plasticity gradient along basal dendrites. (See Figure 2, but now for a layer 2/3 neuron morphology).

**Figure S3.**
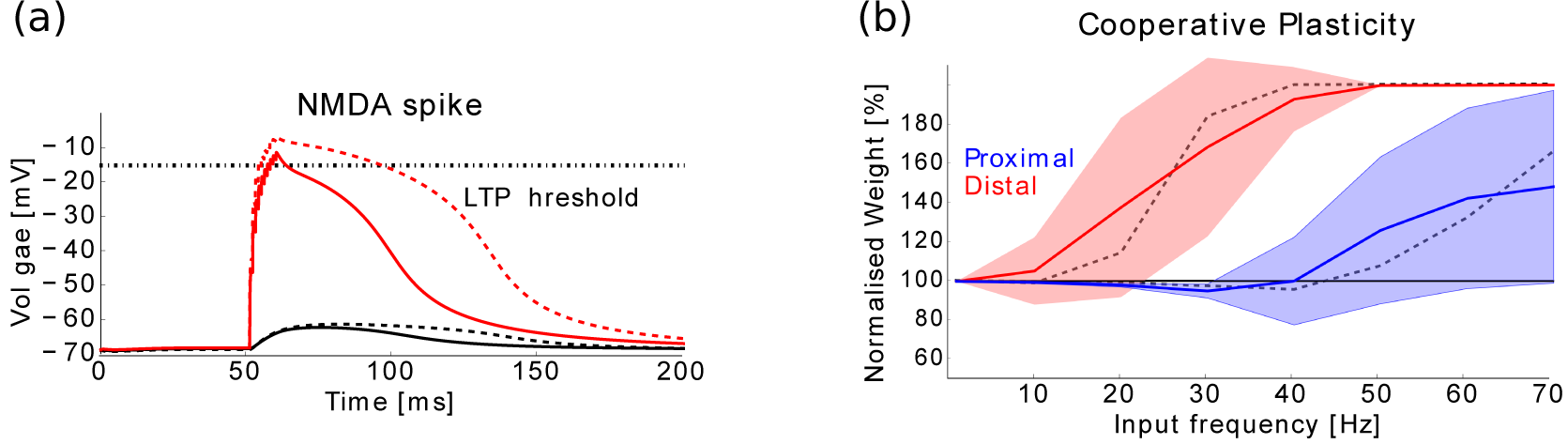
Plasticity of AMPA receptor channels only. (a) When NMDA receptor channels are not plastic, the initial NMDA to AMPA ratio changes due to plasticity. In our model, when starting with an initial weight at half of the maximum and with an initial 2:1 NMDA/AMPA ratio, the latter becomes a 1 to 1 ratio when the AMPA channels reach the maximum bound. Following the same activations as in Figure 2e, the full lines represent an NMDA spike evoked by synapses at the maximum weight and with 1:1 NMDA/AMPA ratio, while the dotted lines represent the same situation with a 2:1 ratio (as in Figure 2e). Even though the NMDA component is weaker in the former situation, an NMDA spike and hence dLTP is still evoked, but less strongly. (b) The same protocol as in Figure 2f leads to similar results (dotted line shows the original mean values).

**Figure S4.**
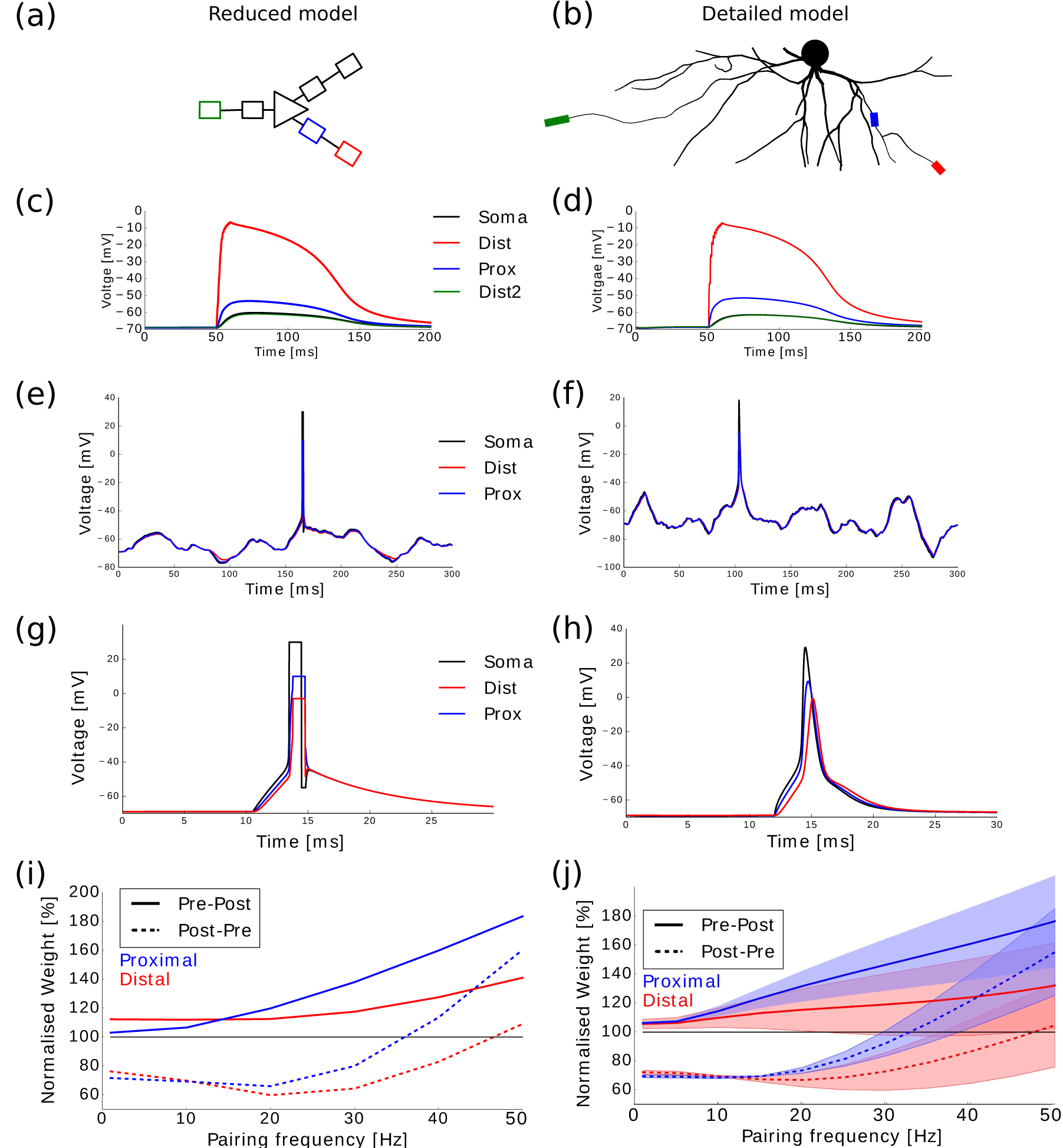
Comparison of the reduced and the detailed model. (a,b) Dendrites in the reduced model are composed of two compartments, one representing a proximal region and one a distal region of the full model. (c,d) Evoking an NMDA spike at a distal compartment (red) will propagate a proximal compartment on the same dendrite (blue), the soma (black) and eventually up to distal compartments in other dendrites (green). (e,f) Noisy currents are injected in the soma of both models. Somatic (black), proximal (blue) and distal (red) voltages are shown. (g,h) Comparison of the spikes in both models. While in the reduced model all spikes have identical shapes, in the full model the spike can vary somewhat depending on the input strength and firing rate. (i,j) Comparison of the plasticity protocol as in Figure 2b gives similar results.

